# A potent human monoclonal antibody with pan-neutralizing activities directly dislocates S trimer of SARS-CoV-2 through binding both up and down forms of RBD

**DOI:** 10.1101/2021.11.29.470356

**Authors:** Xiaofei Wang, Ao Hu, Xiangyu Chen, Yixin Zhang, Fei Yu, Shuai Yue, Arong Li, Junsong Zhang, Zhiwei Pan, Yang Yang, Yao Lin, Leiqiong Gao, Jing Zhou, Jing Zhao, Fang Li, Yaling Shi, Feng Huang, Xiaofan Yang, Yi Peng, Luoyang Tu, Huan Zhang, Huanying Zheng, Jun He, Hui Zhang, Lifan Xu, QizhAo Huang, Yongqun Zhu, Kai Deng, Lilin Ye

**Affiliations:** Institute of Human Virology, Key Laboratory of Tropical Disease Control of Ministry of Education, Zhongshan School of Medicine, Sun Yat-sen University, Guangzhou, Guangdong, China; Department of Gastroenterology of the Second Affiliated Hospital School of Medicine, and Life Sciences Institute, Zhejiang University, Hangzhou, Zhejiang, China; The MOE Key Laboratory for Biosystems Homeostasis & Protection and Zhejiang Provincial Key Laboratory of Cancer Molecular Cell Biology, Life Sciences Institute, Zhejiang University, Hangzhou, Zhejiang, China; School of Laboratory Medicine and Biotechnology, Southern Medical University, Guangzhou, Guangdong, China; Institute of Cancer, Xinqiao Hospital, Third Military Medical University, Chongqing 400038, China; Medical Research Center, Guangdong Provincial People’s Hospital, Guangdong Academy of Medical Sciences, Guangzhou, Guangdong, China; Institute of Immunology, PLA, Third Military Medical University, Chongqing, China; Biomedical Analysis Center, Third Military Medical University, Chongqing, China; Guangzhou Eighth People’s Hospital, Guangzhou Medical University, Guangzhou, Guangdong, China; Guangzhou Women and Children Medical Center, Guangzhou Medical University, Guangzhou, Guangdong, China; Guangdong Provincial Center for Disease Control and Prevention, Guangzhou, Guangdong, China; Center for Cell Lineage and Development, Guangzhou Institutes of Biomedicine and Health, Chinese Academy of Sciences, Guangzhou, Guangdong, China

**Author notes:** Correspondence: Yongqun Zhu, Kai Deng and Lilin Ye. These authors contributed equally to this work (Xiaofei Wang, Ao Hu, Xiangyu Chen, Yixin Zhang, Fei Yu).

## Abstract

The severe acute respiratory syndrome coronavirus 2 (SARS-CoV-2) has caused a global pandemic of novel corona virus disease (COVID-19). The neutralizing monoclonal antibodies (mAbs) targeting the receptor binding domain (RBD) of SARS-CoV-2 are among the most promising strategies to prevent and treat COVID-19. However, SARS-CoV-2 variants of concern (VOCs) profoundly reduced the efficacies of most of mAbs and vaccines approved for clinical use. Herein, we demonstrated mAb 35B5 efficiently neutralizes both wild-type (WT) SARS-CoV-2 and VOCs, including B.1.617.2 (delta) variant, *in vitro* and *in vivo*. Cryo-electron microscopy (cryo-EM) revealed that 35B5 neutralizes SARS-CoV-2 by targeting a unique epitope that avoids the prevailing mutation sites on RBD identified in circulating VOCs, providing the molecular basis for its pan-neutralizing efficacy. The 35B5-binding epitope could also be exploited for the rational design of a universal SARS-CoV-2 vaccine.

## Introduction

As of October 26^th^, 2021, the novel coronavirus SARS-CoV-2 has resulted in more than 243.24 million cases and 4.94 million fatalities^1^. Though an unprecedentedly large number of vaccines and neutralizing mAbs have been developed to contain COVID-19 in the past year, a major concern is the emergence of more transmissible and/or more immune evasive SARS-CoV-2 VOCs, which are antigenically distinct and become dominant in the COVID-19 prevalence over time^2, 3^. Indeed, the D614G variant became prevalent in the early phase of the pandemic and was associated with higher transmission rate ^4^. As the thriving pandemic continued, a rapid accumulation of mutations was observed in SARS-CoV-2 and thus seeded the simultaneous appearance of a plethora of VOCs, which include but not limited to B.1.1.7 (UK; alpha variant)^5^, B.1.351 (SA; beta variant)^6^, P.1 (Brazil; gamma variant)^7^ and B.1.617.2 (India; delta variant)^8^.

In the RBD of SARS-CoV-2 spike protein, B.1.1.7 harbors a N501Y mutation and thus acquires enhanced binding of RBD to the human receptor ACE2^3, 5^. Along with the N501Y mutation, B.1.351 and P.1 develop additional K417N/T and E484K mutations^6, 7^. Meanwhile, B.1.617.2 carries E484Q/L452R mutations^8^. These mutations contribute to the immune escape of SARS-CoV-2 VOCs against many mAbs^3, 9,10,11^, including those already approved for clinical use (casirivimab, bamlanivimab, regdanvimab). These mutant VOCs also undermine humoral immune response elicited by the WT SARS-CoV-2 infection or vaccines targeting WT SARS-CoV-2 protein sequence^10,11,12,13,14,15^. Thus, highly potent and broadly neutralizing mAbs targeting multiple SARS-CoV-2 VOCs are urgently needed for emergency use and elucidating the underlying neutralizing mechanisms of broadly neutralizing mAbs will also provide important insights into the rational design of universal SARS-CoV-2 vaccines.

## Results

### Isolation and characteristics of mAbs 35B5 and 32C7

To discover potent broadly neutralizing mAbs against circulating SARS-CoV-2 VOCs, we adapted a pipeline to rapidly isolate and characterize mAbs (Supplementary fig. 1a). Given the vigorous SARS-CoV-2-specific memory B cell response in individuals recovering from severe COVID-19 illness^16, 17^, cryopreserved PBMCs from these convalescent patients with WT SARS-CoV-2 infection were stained for memory B cell markers (CD19, CD20 and IgG) and avidin-tagged biotinylated SARS-CoV-2 RBD antigen bait. As expected, we found SARS-CoV-2 RBD-specific memory B cells only enriched in PBMCs of convalescent COVID-19 patients, but not healthy donors (Supplementary fig. 1b). Each individual of SARS-CoV-2 RBD-specific memory B cells was further sorted to clone heavy and light chain pairs for mAb production. Two mAbs potentially with superior neutralization activity, one VH3-9/VK2-28 mAb named as 35B5, characterized by a heavy chain CDR3 region of 23 residues and a light chain CDR3 region of 9 residues and another VH3-30/VK4-1 mAb named as 32C7, featured by heavy chain CDR3 of 16 residues and light chain CDR3 of 9 residues (Supplementary fig. 1c, d), were cloned, expressed and analyzed.

Real-time association and dissociation of 35B5 and 32C7 binding to the RBD of SARS-CoV-2 virus were monitored using the surface plasmon resonance (SPR)-based optical assay. We found that 35B5 and 32C7 exhibited different binding affinity in binding to SARS-CoV-2 RBD. Notably, 35B5 exhibited fast-on/slow-off kinetics with an equilibrium dissociation constant (K_D_) of 2.19×10^-12^ M and its binding affinity for SARS-CoV-2 RBD is close to the detection limit at sub-nM level (Fig. 1a); whereas 32C7 showed slow-on/fast-off kinetics with a K_D_ of 1.09×10^-8^ M (Fig. 1b). Consistently, these binding modes were further evidenced by enzyme-linked immunosorbent assay (ELISA) for SARS-CoV-2 RBD, with a EC_50_ value of 0.0183 µg/ml for 35B5 and a EC_50_ value of 0.1038 µg/ml for 32C7 (Supplementary fig. 1e).

**Fig. 1.**
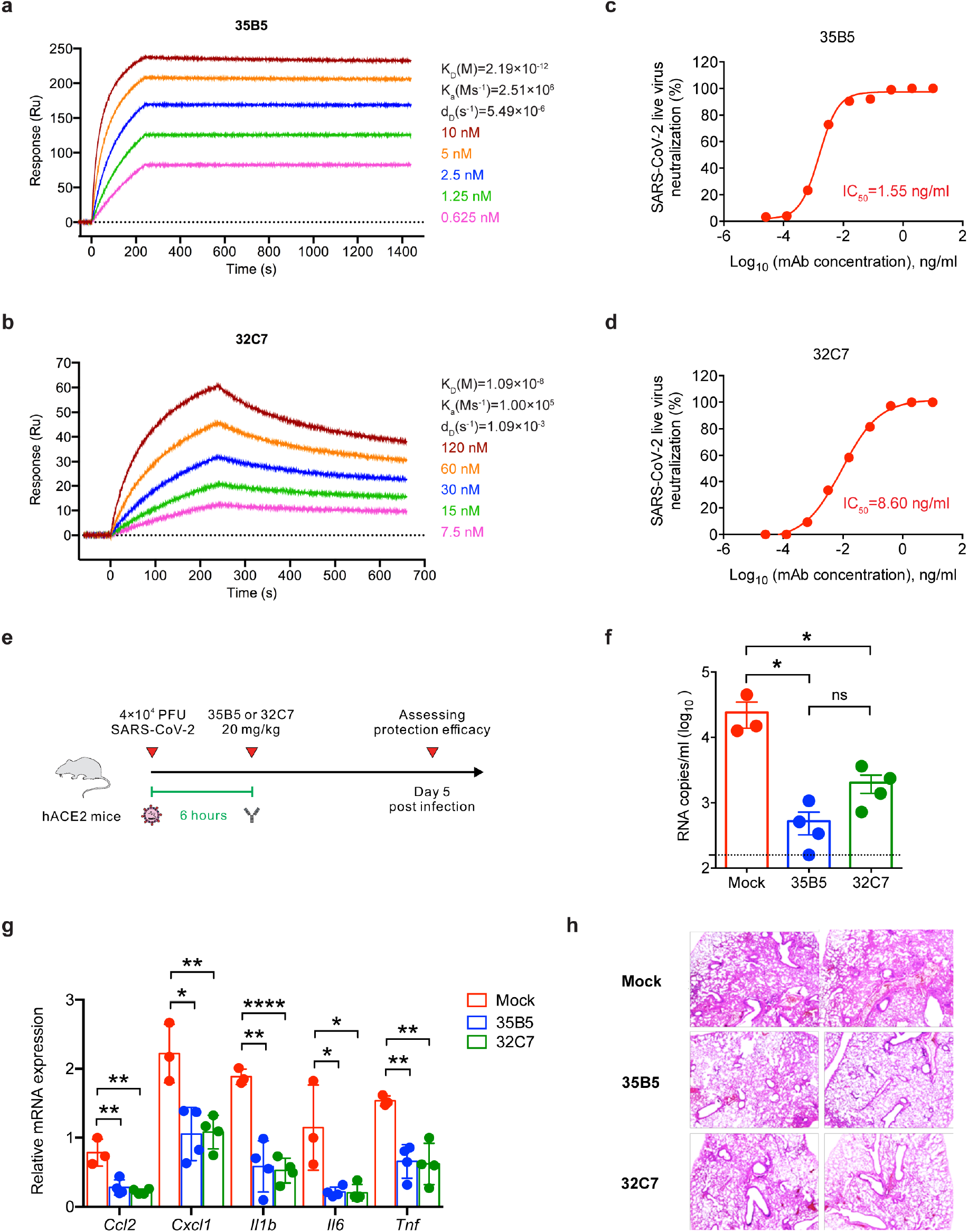
35B5 and 32C7 protect against authentic SARS-CoV-2 virus. **a**, **b** Affinity analysis of 35B5 (**a**) and 32C7 (**b**) binding to immobilized SARS-CoV-2 RBD by using biolayer interferometry. **c, d** *In vitro* neutralizing activity of 35B5 (**c**) and 32C7 (**d**) against authentic SARS-CoV-2. Mixture of SARS-CoV-2 and serially diluted 35B5 or 32C7 were added to Vero E6 cells. After 48 hours, IC_50_ values were calculated by fitting the viral RNA copies from serially diluted mAb to a sigmoidal dose-response curve. **e** Schematic diagram of 35B5 and 32C7 treatment *in vivo*. Six hours after infection with 4×10^4^ PFU SARS-CoV-2, the hACE2 mice received a single dose of 35B5 or 32C7 with 20 mg/kg or no mAb treatment (mock). At day 5 post infection, lung tissues were collected for viral burden assessment, cytokine/chemokine assay and histological analysis. **f** Viral titers in the lungs were measured by qRT-PCR and presented as RNA copies per milliliter of lung abrasive fluid. **g** Gene expressions of cytokines and chemokines in the lungs were determined by qPT-PCR. **h** Histopathological analysis of lung tissues. The data are representative of at least two independent experiments. \**P* < 0.05, \*\**P* < 0.01 and \*\*\*\**P* < 0.0001. Not significant, ns. Error bars in (**f**) and (**g**) indicate SD.

### Potent neutralization capacity of mAbs 35B5 to SARS-CoV-2 virus *in vitro* and *in vivo*

We next investigated the neutralizing capacity of 35B5 and 32C7 against authentic WT SARS-CoV-2 infection in Vero E6 cells. Remarkably, we found that both 35B5 and 32C7 neutralized authentic SARS-CoV-2 virus in the low picomolar range, with observed IC_50_ values of 1.55 ng/ml for 35B5 (Fig. 1c) and 8.60 ng/ml for 32C7 (Fig. 1d). We further assessed *in vivo* protection efficacy of 35B5 and 32C7 in the human ACE2 (hACE2)-expressing transgenic mouse model (ICR background) that is sensitized to SARS-CoV-2 infection^18^. The hACE2-humanized mice were treated intraperitoneally with a single dose of 35B5 or 32C7 with 20 mg/kg at 6 hours after intranasal infection with 4×10^4^ PFU of SARS-CoV-2. As control, infected mice of mock group were administrated with an equal volume of PBS (Fig. 1e). At day 5 post-infection, the viral loads in the lungs of the mock group surged to ∼10^4^ RNA copies/ml (Fig. 1f). By contrast, 35B5 and 32C7 treatment remarkably reduced the viral titers, with ∼10^2^ RNA copies/ml resulting 100-fold reduction and ∼10^3^ RNA copies/ml resulting 10-fold reduction, respectively (Fig. 1f). In addition, we also determined whether 35B5 and 32C7 treatment ameliorated the pathological lung damage in the hACE2 mice infected with SARS-CoV-2. The transcripts of cytokines and chemokines (e.g., *Ccl2*, *Cxcl1*, *Il1b*, *Il6* and *Tnf*), indicative of tissue inflammation, were greatly reduced in both 35B5- and 32C7-treated groups when compared to those observed in the mock group (Fig. 1g). Parallelly, the SARS-CoV-2 virus caused interstitial pneumonia characterized by inflammatory cell infiltration, alveolar septal thickening and distinctive vascular system injury in the mock group, but not in 35B5 or 32C7 treatment groups; additionally, less pronounced lesions of alveolar epithelial cells or focal hemorrhage were observed in the lung sections from mice that received 35B5 or 32C7 treatment than in those from mice of mock group (Fig. 1h). Thus, these results together indicate the potential therapeutic role for these two mAbs, in particular 35B5, in treating COVID-19.

### 35B5 exhibits ultrapotent and broad neutralization activity to SARS-CoV-2 VOCs *in vitro* and *in vivo*

We then sought to determine the neutralizing capacity of 35B5 and 32C7 mAbs against SARS-CoV-2 variants. Neutralizing activity of these two mAbs was assessed against authentic SARS-CoV-2 viruses, including WT, D614G, B.1.351 and B.1.617.2 strains. It turned out 35B5 preserved highly potent neutralizing capacity against the D614G variant (IC_50_=7.29 ng/ml), the B.1.351 variant (IC_50_=13.04 ng/ml) and the B.1.617.2 variant (IC_50_=5.63 ng/ml) (Fig. 2a); however, neutralizing capacity of 32C7 against these two variants was severely blunted, with IC_50_ values of 127.60 ng/ml for the D614G variant, 1420.00 ng/ml for the B.1.351 variant and 3442.00 ng/ml for the B.1.617.2 variant (Fig. 2b). In parallel, we also explored the neutralization of authentic virus by 35B5 and 32C7 in a focus forming assay (FFA). Consistently, we found that 35B5 showed potent neutralizing capacity against the WT SARS-CoV-2, the D614G variant, the B.1.351 variant as well as the B.1.617.2 variant (Fig. 2c); whereas 32C7 exhibited a clear reduction in neutralization against the D614G variant, the B.1.351 variant and the B.1.617.2 variant (Fig. 2d). Given the highly broad neutralization activities of 35B5 *in vitro*, we next assessed the *in vivo* protection efficacy of 35B5 against the D614G variant, the B.1.351 variant and the B.1.617.2 variant. To this end, a different strain of hACE2-humanized mice (C57BL/6 background)^19^ were infected with 4×10^4^ PFU of SARS-CoV-2 D614G or B.1.351 or B.1.617.2 and then treated with a single dose of 35B5 with 30 mg/kg or PBS at 4 hours after infection. Viral loads in the lungs of all infected mice were measured at day 5 post-infection (Fig. 2e). Indeed, we found that 35B5 treatment resulted in a ∼10-fold, ∼100-fold and ∼100-fold reduction of viral titers in the lungs of mice infected with D614G, B.1.351 and B.1.617.2, respectively (Fig. 2f). Moreover, tissue inflammation and interstitial pneumonia caused by SARS-CoV-2 D614G/B.1.351/B.1.617.2 infection were largely ameliorated in 35B5-treated groups compared to that in mock groups (Fig. 2g-j). Together, these results suggest that 35B5 potently neutralizes a variety of SARS-CoV-2 variants as a broadly neutralizing mAb.

**Fig. 2.**
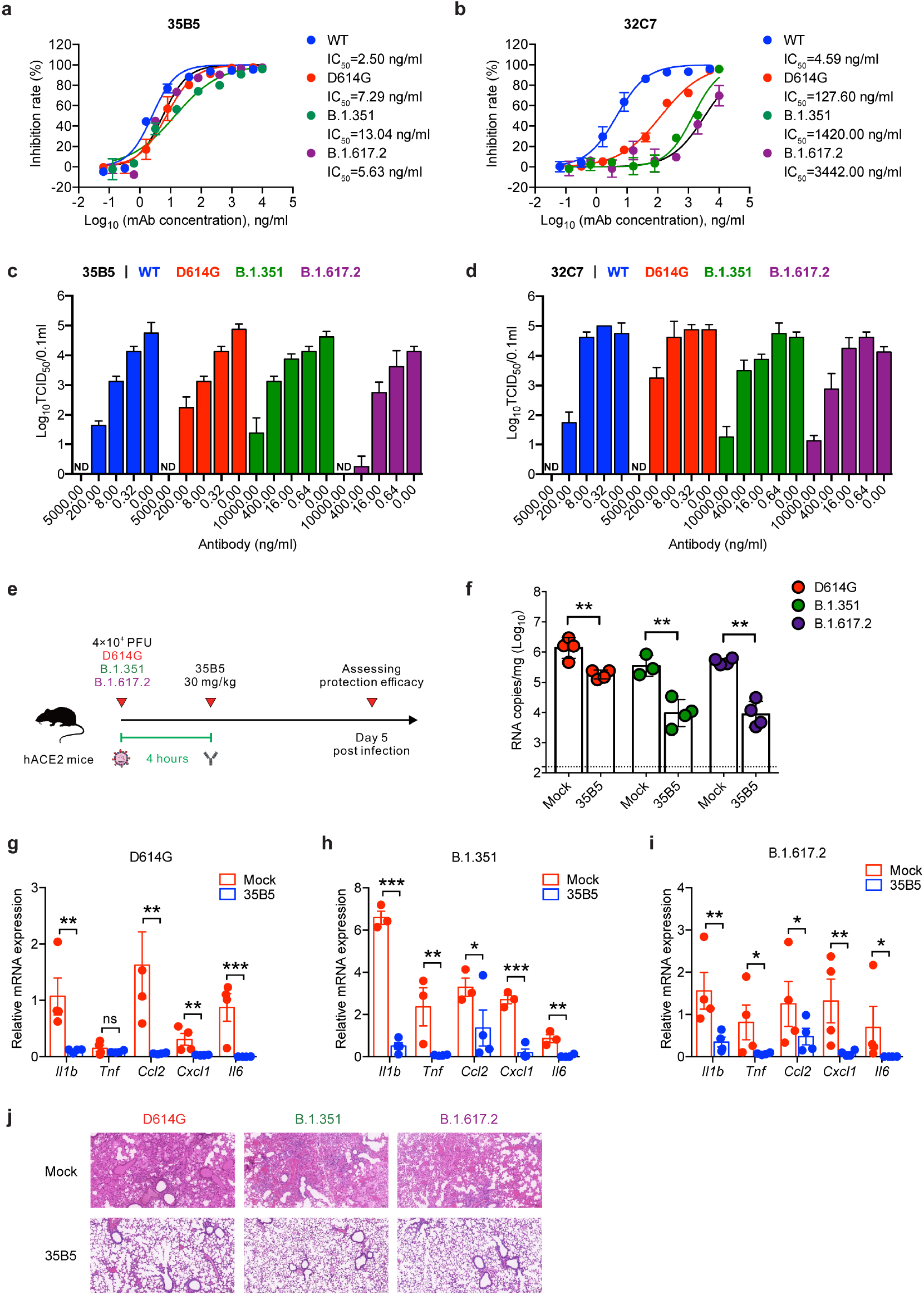
Neutralizing activity of 35B5 against SARS-CoV-2 VOCs. **a, b** Neutralizing activity of 35B5 (**a**) and 32C7 (**b**) against authentic SARS-CoV-2 viruses, including WT strain, D614G variant, B.1.351 variant and B.1.617.2 variant. Mixture of each SARS-CoV-2 strain and serially diluted 35B5 or 32C7 were added to Vero E6 cells. After 48 hours, IC_50_ values were calculated by fitting the viral RNA copies from serially diluted mAb to a sigmoidal dose-response curve. **(c, d)** FFA analysis of 35B5 (**c**) and 32C7 (**d**) against WT SARS-CoV-2, D614G variant, B.1.351 variant and B.1.617.2 variant. Virus cultures were serially diluted and added to Vero E6 cells. After 24 hours, the foci were visualized by using HRP-conjugated polyclonal antibodies targeting SARS-CoV-2 nucleocapsid protein in Vero E6 cells and counted with a ELISPOT reader. **e** Schematic diagram of 35B5 treatment *in vivo*. Four hours after infection with 4×10^4^ PFU SARS-CoV-2 D614G or B.1.351 or B.1.617.2, the hACE2 mice received a single dose of 35B5 with 30 mg/kg or no mAb treatment (mock). At day 5 post infection, lung tissues were collected for viral burden assessment. **f** Viral titers in the lungs were measured by qRT-PCR and presented as RNA copies per gram of lung tissue. **g to i** Gene expressions of cytokines and chemokines in the lungs were determined by qPT-PCR. **j** Histopathological analysis of lung tissues. ND, not detected. Data are representative of two independent experiments. Error bars in **a**-**d**, **f-i** indicate SD.

### Complex structure of mAb 35B5 with the spike protein

To investigate the structural basis of the superior and broad neutralizing activity of 35B5 against the SARS-CoV-2 variants, we employed cryo-EM approaches to determine the complex structure of the Fab region of 35B5 with the spike protein of SARS-CoV-2. Incubation of 35B5 Fab with the ectodomain of the spike protein S-2P^20^, a stabilized spike mutant, severely caused the dissociation of the trimer and disrupted its structure quickly *in vitro* (Supplementary Fig. 2). 35B5 Fab even disrupted the ectodomain trimer structure of the spike S-HexaPro protein (S-6P)^21^, a more stable spike variant containing four additional proline substitutions (F817P, A892P, A899P and A942P) from S-2P (Supplementary Fig. 2), suggesting that 35B5 harbors the potent dissociation activity toward the spike protein. Nonetheless, a few of the S-6P particles were still found to maintain the triangular architecture^21, 22^ after 35B5 Fab treatment for 3 min, which thereby allowed us to carry out cryo-EM analyses of the 35B5 Fab-S-6P complex. We successfully determined the structures of the 35B5 Fab-S-6P complex in three conformational states to the resolutions of 3.7 Å, 3.4 Å and 3.6 Å, respectively (Supplementary Fig. 3 and Supplementary Table 1).

In the 35B5 Fab-S-6P complex structure of State 1, two RBD domains of the S-6P trimer are in the standard “up” conformations^20^ and are bound by 35B5 Fabs (Fig. 3a). The other RBD domain is in the “down” conformation^20^ as that was found in the Fab-free spike trimers. In the 35B5 Fab-S-6P complex structure of State 2, each of the three RBD domains was in the “up” conformation and bound by a 35B5 Fab (Fig. 3a). In the 35B5 Fab-S-6P complex structure of State 3, although all the three RBD domains were bound by 35B5 Fab, only one RBD domain maintained the “up” conformation. The other two RBD domains were in unprecedented conformations, which we for the first time named as “releasing” conformations (Fig. 3a). Compared to the “up” RBDs, the two “releasing” RBD domains move out by 6.4 Å and 23.0 Å, respectively (Fig. 4f). The two “releasing” RBD domains generated large gaps with the adjacent NTD domains in the S-6P trimers (Fig. 3a), suggesting that the spike protein is undergoing structural dissociation.

**Fig. 3.**
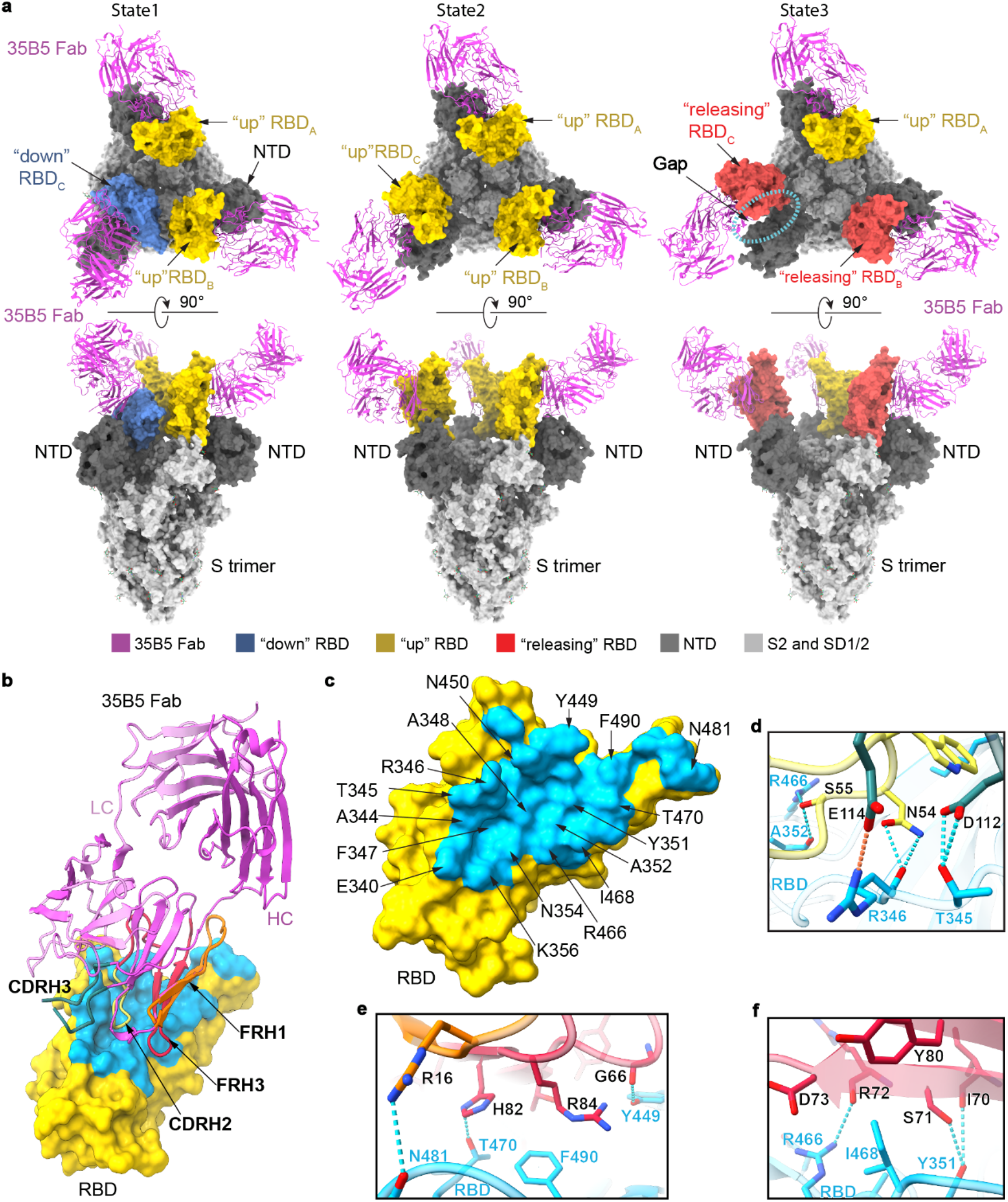
Cryo-EM structures of the spike protein S-6P complexed with 35B5 Fab. **a** The structures of the S-6P-35B5 Fab complex in three states. The S trimer is represented as surface. 35B5 Fab is shown in cartoon and colored in purple. The “down”, “up” and “releasing” RBD domains are colored in blue, yellow and red, respectively. The NTD domain of the S trimer is colored in deep grey. The SD1, SD2 and S2 domains are colored in light grey. The gap caused by 35B5 Fab between the “releasing” RBD and NTD domains is highlighted with dashed lines. **b** Interactions of 35B5 Fab with “up” RBD. The interacting residues within 4 Å in RBD are colored in cyan. The RBD-interacting regions CDRH2, CDRH3, FRH1 and FRH3 of 35B5 Fab are colored in yellow, blue, orange and red, respectively. **c**, The 35B5 epitope on RBD. The epitope residues are labeled as indicated. **d-f** Detailed interactions of the CDR (**d**) and FR regions (**e** and **f**) of 35B5 Fab with RBD. Hydrogen-bond and salt-bridge interactions are shown as blue and orange dashed lines, respectively.

**Fig. 4.**
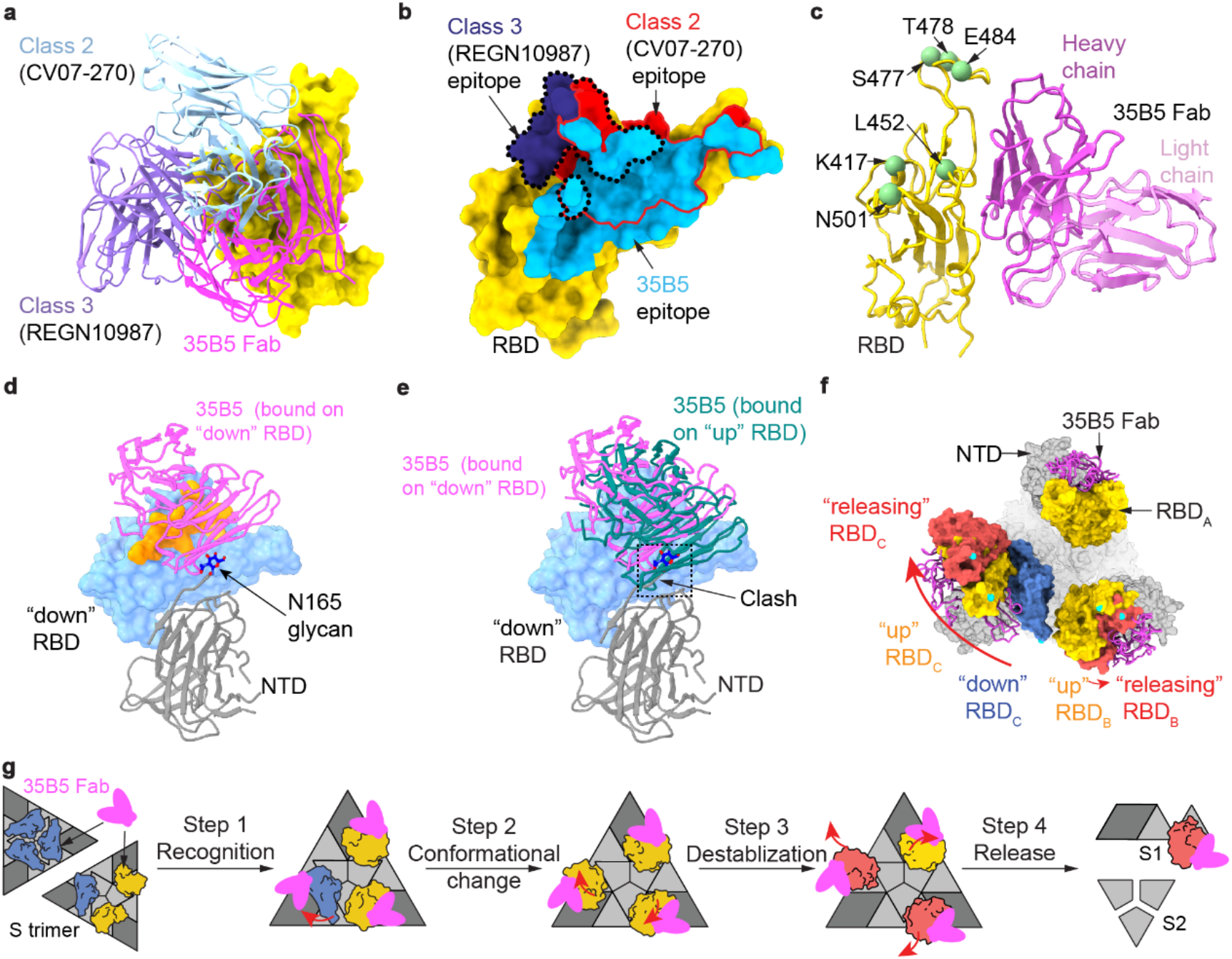
Structural basis and neutralizing mechanism for the broad neutralizing activities of 35B5 to the SARS-CoV-2 variants. **a** Structural comparisons of the 35B5 Fab-“up” RBD interactions with those of the previously identified Class 2 and 3 neutralizing antibodies. 35B5 Fab and the Fab regions of the representative antibodies CV07-270 and REGN10987 from Class 2 and 3, respectively, are illustrated in cartoon and colored as indicated. RBD is represented as surface in yellow. **b** Comparison of the epitope of 35B5 with those of the representative neutralizing antibodies of Class 2 and 3 on RBD. The 35B5 epitope is colored in cyan. The epitopes of CV07-270 (Class 2) and REGN10987 (Class 3) are labeled using red and black lines, respectively. **c** Mapping of the frequently mutated residues of the spike protein in the recent SARS-CoV-2 variant in the 35B5 Fab-RBD structure. RBD and 35B5 Fab are shown in cartoon and colored in yellow and purple, respectively. The frequently mutated residues in RBD in the recent SARS-CoV-2 variant are highlighted with green spheres. **d** Interactions of 35B5 Fab with the “down” RBD. 35B5 Fab, the “down” RBD and NTD domains are colored in purple, cyan and grey, respectively. The 35B5 Fab-interacting region on RBD is highlighted in orange. **e** Structural clashes between 35B5 Fab and the NTD domain upon structural superimposition of the “up” RBD-35B5 Fab model with the “down” RBD-35B5 Fab region in the state 1 S-6P-35B5 Fab complex. The structural clashes are highlighted with dashed lines. **f** Conformational changes of the RBD domains upon 35B5 Fab binding. The S-6P-35B5 Fab complex structures of three states were superimposed. 35B5 Fabs are shown in cartoon in purple. The “down”, “up”, and “releasing” RBD domains are colored in blue, yellow and red, respectively. The conformational changes of RBDs are indicated with arrows. **g** Schematic diagram of the neutralizing mechanism of 35B5 against SARS-CoV-2. The “down” and “up” RBDs in the tight-closed or loose-closed S trimers are colored in blue and yellow, respectively. The conformation changes and transmission of RBDs from the “down” to “up”, finally to “releasing” states are shown as red arrows.

### 35B5 targets a unique epitope for pan-neutralizing activity

The interactions of 35B5 Fab with the “up” RBDs are identical to those of 35B5 Fab with the “releasing” RBDs in the three states of the 35B5 Fab-S-6P complex. The interface covers a largely buried area of ∼1029 Å^2^ (Fig. 3b, c). The epitope in RBD for 35B5 is composed of 30 interacting residues including R346, F347, N354, R466, A352, K444, Y449, N450, R466, I468, T470, N481, and F490, which form extensive hydrophilic interactions with 35B5 Fab in the structure (Fig. 3b-f). The corresponding paratope in 35B5 Fab consists of two heavy chain complementarity determining regions (CDRH2 and CDRH3) and the heavy chain frameworks (FRH1 and FRH3). The epitope of 35B5 on RBD is distinct from those of the previously identified 4 classes of neutralizing antibodies to RBD^23^. The 35B5 Fab-binding surface on RBD is located at the site opposite to the receptor ACE2-binding surface^23,24,25,26^, which is targeted by the class 1 antibodies, suggesting that 35B5 doesn’t directly block the receptor recognition for neutralization. Although the epitope of 35B5 Fab involves some regions of the epitopes of the classes 2 and 3 of antibodies (Fig. 4a, b and Supplementary Fig. 4a), the major 35B5-interacting residues, including the SARS-CoV-2 specific residues N354, T470 and N481 (Fig. 3b), which are not conserved in the spike proteins of SARS-CoV and MERS-CoV, are outside of the epitopes of the classes 2 and 3 of antibodies. Therefore, 35B5 targets a distinctive epitope to specifically neutralize SARS-CoV-2.

The SARS-CoV-2 VOCs contains several prevailing mutations on RBD, including N501Y (B.1.1.7 (alpha), B.1.351 (beta) and P1 (gamma)), K417N (B.1.351 (beta) and P1 (gamma)), L452Q (C.37 (lambda)), L452R (B.1.617.2 (delta), B.1.427/B.1.429 (epsilon), B.1.617.1 (kappa) and B.1.526 (iota)), S477N (B.1.526 (iota)), T478K (B.1.617.2 (delta)), E484K (B.1.351 (beta), P1 (gamma), B.1.617.2 (delta), B.1.525 (eta), and B.1.526 (iota)) and F490S (C.37 (lambda)) (Fig. 5a). However, in the 35B5 Fab-S-6P complex structures, the residues N501, K417, L452, S477, T478 and F490 are not involved in the 35B5-RBD interactions. Only the residue E484 is located at the edge of the 35B5-RBD interface but not contacted by 35B5 Fab (Fig. 4c and Fig. 5b). Substitution of E484 by a lysine residue doesn’t generate severe structural collision with 35B5 Fab (Fig. 5c). It has been found that L452R mutation is of significant adaptive value to the B.1.617.2 variant (delta). However, L452 has the distance of more than 4.5 Å and 5.1 Å, respectively, to the residues T69 and Y60 of 35B5 Fab in the 35B5 Fab-S-6P complex structure (Fig. 5e), suggesting that the residue is not contacted by 35B5 Fab. Substitution for L452 by an arginine does not spatially affect the 35B5 Fab-RBD contacts. Instead, the mutation generates direct interactions with the residues Y60 and T69 of 35B5 Fab within 3 Å and likely forms two hydrogen bonds (Fig. 5e), suggesting that this mutation does not affect the binding affinity of 35B5 to RBD. Consistently, 35B5 has the comparable super-potent neutralization efficacy to the B.1.617.2 variant as that to the wild-type virus (Fig. 2). Recently, it was found that the C.37 variant (lambda) contains the mutations L452Q and F490S^27^. However, the mutations L452Q and F490S would not structurally affect the interactions of 35B5 Fab with RBD in the 35B5 Fab-S-6P complex structures (Fig. 5f, g), indicating that 35B5 might also exhibit potent neutralizing efficacy to the C.37 variant. Thus, the unique epitope of 35B5 on RBD subtly avoids the prevailing mutation sites, which provides the molecular basis for the potent pan-neutralizing efficacy of 35B5 to the SARS-CoV-2 VOCs.

**Fig. 5.**
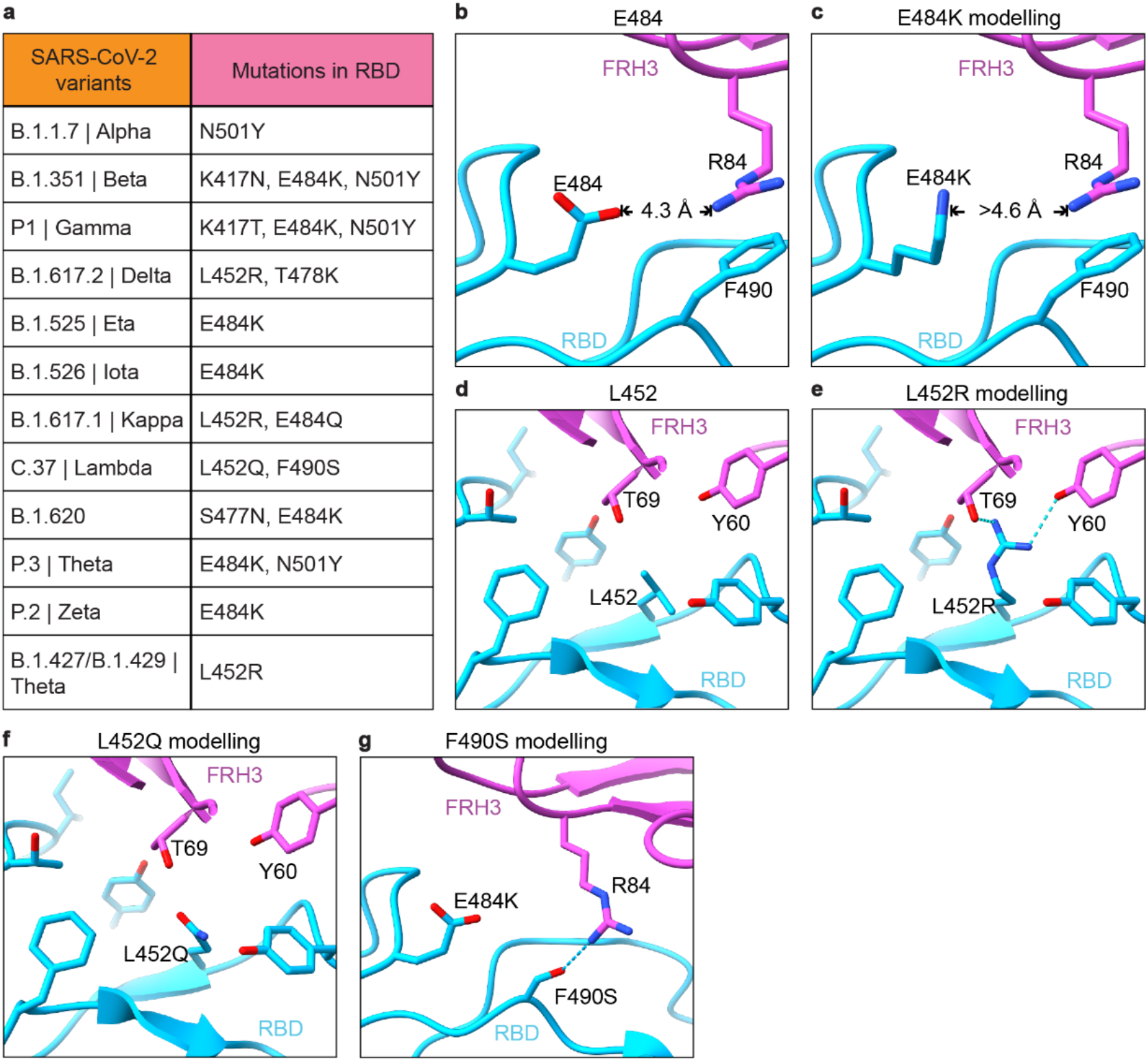
Interactions of 35B5 Fab with RBD. **a** Mutations in the RBD of reported SARS-CoV-2 variants of concern or interest. **b**-**g** The residues E484 (**b**) and L452 (**d**) are not involved in the interactions of RBD with 35B5 Fab. E484 is more than 4.2 Å away from 35B5 R84, which forms cation-π interaction with RBD F490. The E484K mutation in RBD does not interact with 35B5 R84 through molecular modelling (**c**). The L452Q modelling suggests none interactions formed at this site with 35B5 (**f**). In structural modeling, the L452R mutation (**e**) and F490S mutation (**g**) in RBD generates potential additional interactions with 35B5 Fab. Potential hydrogen bonds are shown as dashed lines.

### Neutralization mechanism of mAb 35B5

To investigate the neutralization mechanism of 35B5, we analyzed the “down” RBD domain in the State 1 35B5 Fab-S-6P complex. In the density map of the State 1 complex, there were some residual densities closed to the “down” RBD domain. We further carried out local refinements on the “down” RBD. The local refinements generated a 4.8-Å density map and revealed that there is a 35B5 Fab contacting the edge of the epitope on the “down” RBD. The low resolution of the local refinement map suggests that the interaction between the 35B5 Fab and the “down” RBD is highly dynamic (Supplementary Fig. 4b-e). Structural modeling after the local refinements revealed that, in contrast to the neutralizing mAbs BD-368^21, 22^ and C002^23^ recognizing the epitopes that are fully exposed in both “down” and “up” RBDs, 35B5 Fab utilizes the CDRH regions to interact with the residues E340, T345, R346, F347, R346, K444, Y449 and N450 at the exposed edge of the epitope on the “down” RBD, indicating that this RBD domain is being initially recognized by 35B5 (Fig. 4d and Supplementary Fig. 4e and Supplementary Fig. 7a, c). Structural superimposition of the 35B5 Fab-“up” RBD model with the “down” RBD in the 35B5 Fab-S-6P complex of State 1 or those in the Fab-free spike trimers^21, 28^ reveals that the β-sheet and the linking loop of FRH1 of 35B5 Fab have severe structural clashes with the adjacent NTD domain of the spike protein (Fig. 4e), suggesting that upon the high-affinity binding of 35B5 Fab onto the “down” RBD, the spatial collisions between 35B5 Fab and the NTD domain potentially exert repulsion force onto the NTD domain to induce the conformational conversion of RBD from “down” to “up” conformation. Further outward movement of the “up” RBDs generated the “releasing” conformations in the State 3 35B5 Fab-S-6P complex. Thus, the 35B5 Fab-S-6P complex structures in these three states suggest that neutralization of SARS-CoV-2 by 35B5 is likely carried out in four sequential steps (Fig. 4g): 35B5 firstly binds to the exposed edge of the epitope to recognizes the “down” RBDs of the spike protein; subsequently, binding of 35B5 imposes structural clashes on the NTD domain to drive the conformational changes of RBDs from “down” to “up”; next, the unstable “up” conformations of the RBD domains destabilizes the structure of the spike trimer and induces the outward movement and releasing of RBDs; finally, the released RBD domains cause the dissociation of the spike trimer.

The SARS-CoV-2 VOCs contains the most prevalent mutation D614G in the spike protein, which enhances infectivity by inducing the wedge of a disordered loop between the SD1/CTD1 and NTD domains within a protomer to prevent premature dissociation of the G614 trimer^29^. The residue D614 is not located in the epitope of 35B5 (Fig. 3b), thereby having no effects on its binding to the RBD domain. Moreover, upon binding to the “down” RBD domain, the capability of 35B5 to exert the repulsion force onto the NTD domain can counteract the effects of the D614G mutation on the structural arrangement of the spike protein. Therefore, the unique epitope of 35B5 avoiding the prevailing mutation sites on RBD and the repulsion force of 35B5 exerting onto NTD during initial recognition renders its super-potent pan-neutralizing efficacy to the SARS-CoV-2 VOCs.

### Cryo-EM structure of mAb 32C7

In contrast to 35B5, mAb 32C7 could not efficiently neutralize the SARS-CoV-2 VOCs. We also solved the cryo-EM structure of the Fab region of mAb 32C7 (hereafter named as 32C7 Fab) in complex with S-6P at a resolution of 2.8 Å (Supplementary Fig. 5 and Supplementary Table 1). The complex structure only contains one 32C7 Fab molecule, which is bound to a “down” RBD domain in the spike protein (Supplementary Fig. 6a-f). The binding surface of 32C7 Fab on RBD overlaps with the epitopes of the class 3 antibodies (Supplementary Fig. 6c), suggesting that 32C7 belongs to the classic class 3 family of neutralizing antibodies. The 32C7 Fab-RBD interface only covers a buried area of approximately 935 Å^2^, which is much less than that of 35B5 Fab (Fig. 3c and Supplementary Fig. 6g and Supplementary Fig. 7b, d). The smaller epitope surface on RBD determined the weaker neutralization efficacy of 32C7 as compared to that of 35B5 (Fig. 1). In the S-6P-32C7 Fab complex structure, 32C7 Fab does not structurally clash with the NTD domain. Instead, 32C7 Fab interacts with the glycan moiety on the residue N165 of the NTD domain (Supplementary Fig. 6b). In addition to the D614G mutation, the B.1.351 variant harbors 4 amino acid substitutions and 1 deletion of 3 amino acids, which cause the conformational change of the NTD domain. The conformational changes of the NTD domain potentially weaken the interactions of 32C7 Fab with the glycan moiety of N165, and further affect its epitope recognition of RBD, thereby impairing the neutralizing efficacy of 32C7 Fab against the B.1.351 and other VOCs.

## Discussion

In this study, we demonstrate 35B5 as a ultrapotent and pan-neutralizing human monoclonal antibody against currently circulating SARS-CoV-2 provided by experimental evidence *in vitro* and *in vivo* as well as structural analysis. By contrast, many mAbs such as casirivimab (REGN10933), bamlanivimab (LY-CoV555), etesevimab (LY-CoV016), regdanvimab (CT-P59), ABBV-2B04 (2B04) and 32C7 in the study partially or entirely lose the neutralizing activity against B.1.351 and B.1.617.2^2, 10, 30, 31^. The mechanisms underlying the picomolar broad neutralization by 35B5 at least involve three aspects: 1) broad interface and extensive interactions between 35B5 and RBD endow 35B5 as a potent cross-epitope mAb; 2) proactive dissociation of the spike trimer by structural clashes between 35B5 Fab FRH1 and spike NTD domain; 3) no direct contacts between 35B5 Fab and prevailing mutations of SARS-CoV-2 VOCs.

Previous works indicated that the ACE-binding surface is partially exposed on the “down” RBDs in the tight- or loose-closed spike trimer^21, 28^. The epitopes of some neutralizing mAbs are also partially exposed on the “down” RBDs^23, 32,33,34,35^. The 35B5 Fab-S-6P complex structures in the three states we determined provide for the first time the direct structural evidence for the possibility that the ACE2 or mAbs can approach the partially exposed surface or epitope residues for initial recognition and fulfil the conformational transformation of RBD. In previously identified RBD-targeted mAbs, almost all class 1 mAbs interact extensively with the residues K417 and N501. Most class 2 and class 3 mAbs contact E484, and most class 3 mAbs interact with L452^3^. 35B5 does not directly bind to these prevailing mutant sites. In contrast to the Class 3 mAb S309^26^ and the Class 4 mAb CR3022^36^, which target the residues conserved in the spike proteins of SARS-CoV, SARS-CoV-2 and MERS-CoV and therefore have cross-species neutralizing activities, 35B5 is a SARS-CoV-2 specific antibody with super potent pan-neutralizing activities to SARS-CoV-2 VOCs.

Protective vaccines are considered as keys to terminate the COVID-19 pandemic. However, accumulating evidence suggests that SARS-CoV-2 VOCs compromise neutralization by antibody responses induced by multiple vaccines^12,13,14,15, 37,38,39^, including mRNA vaccines (BNT162b2 and mRNA-1273), adenovirus-based vaccines (AZD1222 and JNJ-78436735), a nanoparticle-based vaccine (NVX-CoV2373) and an inactivated protein vaccine (Coronavac), primarily due to RBD mutations. Thus, vaccines aimed at eliciting broadly neutralizing mAbs against SARS-CoV-2 VOCs are urgently needed. In the scenario of coping with diverse HIV-1 variants, an experimental pan-HIV-1 vaccine was designed based on the structurally defined epitope of a HIV-1 neutralizing mAb N123-VRC34.01, which targets a linear peptide of fusion peptide that is across-clade conserved^40, 41^. In our study, cryo-EM structure of neutralizing 35B5 to the SARS-CoV-2 spike revealed an epitope that is composed of 30 residues and conserved among all the currently known SARS-CoV-2 VOCs.

Importantly, mAb 35B5 was cloned from SARS-CoV-2 RBD-specific B cells obtained from individuals during their early convalescence, which might endow mAb 35B5 with limited somatic hypermutation (SHM). In contrast, a broad neutralizing mAb, ADG-2, was engineered by *in vitro* affinity optimization^42^ and the extensive SHM makes ADG-2 epitope challenging as a target for SARS-CoV-2 vaccines. Thus, we propose that the 35B5 epitope might be a valuable target for the rational design of a universal SARS-CoV-2 vaccine, which is presumably essential for the ultimate termination of COVID-19 pandemic in the future.

## Methods

### Human samples

The COVID-19 patients enrolled in the study were admitted to Guangzhou Eighth People’s Hospital (January to March 2020) and provided written informed consent. Blood samples were collected from patients during their convalescence and the time between symptom onset to sample collection was around 20 days. Healthy donors were 2 adult participants in the study. Blood samples were collected in cell preparation tubes with sodium citrate (BD Bioscience). Then, peripheral blood mononuclear cells (PBMCs) were isolated from blood samples using Ficoll (TBD Science), washed with PBS, suspended in cell freezing medium (90% FBS plus 10% DMSO), frozen in freezing chamber at -80 °C, and then transferred to liquid nitrogen. The study received IRB approvals at Guangzhou Eighth People’s Hospital (KE202001134).

### Single-cell sorting, RT-PCR and PCR cloning

PBMCs were firstly incubated with Human TruStain FcX (Biolegend) at 4 °C for 30 min and then stained with biotin-conjugated SARS-CoV-2 RBD protein (Sino Biological, 40592-V05H) at 4 °C for 20 min. Next, PBMCs were stained with PE-Cy7-conjugated streptavidin (eBioscience), FITC-conjugated anti-CD19 antibody (Biolegend), PE-conjugated anti-CD20 antibody (Biolegend), APC-conjugated anti-human IgG (Fc) (Biolegend), APC-Cy7-conjugated anti-CD3 antibody (Biolegend), APC-Cy7-conjugated anti-CD14 antibody (Biolegend), APC-Cy7-conjugated anti-CD56 antibody (Biolegend) and APC-Cy7-conjugated LIVE/DEAD dye (Life Technologies) at 4 °C for 30 min. All the stainings were performed in PBS containing 5% mouse serum (wt/vol). For cell sorting, the SARS-CoV-2 RBD-specific IgG^+^ B cells were sorted into 96-well plates, with a single cell and 10 µl catch buffer per well. These plates were stored at -80 °C for further usage. Catch buffer: to 1 ml of RNAase-free water (Tiangen Biotech), add 40 µl Rnasin (NEB) and 50 µl 1.5 M Tris pH 8.8 (Beijing Dingguo Changsheng Biotech).

IgG VH and VL genes from B cells were PCR amplified and cloned into vectors as previously described^43, 44^. Briefly, plates containing cell lysate were thawed on ice, added with RT-PCR master mix and IgG VH/VL primers per well and performed with RT-PCR following the one step RT-PCR kit protocol (Takara, RR057A). Then, RT-PCR products were nested PCR-amplified with primers for IgG VH or IgG VL following the HS DNA polymerase kit protocol (Takara, TAK R010). The IgG VH and IgG VL nested PCR products were purified and next cloned into human IgG1 heavy chain and light chain expression vectors, respectively.

### Monoclonal antibody production and purification

Monoclonal antibodies were produced by using the ExpiCHO^TM^ Expression System (Thermo Fisher). Briefly, ExpiCHO-S^TM^ cells of 200 ml (6×10^6^ cells/ml) in 1-L flask were transfected with master mixture containing 200 µg heavy chain plasmid, 200 µg light chain plasmid and 640 µl ExpiFectamine^TM^ CHO reagent. On the day after transfection, cultured cells were added with 1.2 ml ExpiCHO^TM^ enhancer and 48 ml ExpiCHO^TM^ feed. A second volume of ExpiCHO^TM^ feed was added into cultured cells on day 5 post-transfection. Supernatants were harvested on day 12 post-transfection for antibody purification using rProtein A Sepharose affinity chromatography (Sigma).

### Cells and viruses

Vero E6 cells were obtained from ATCC and maintained in Dulbecco’s Modified Essential Medium (DMEM) supplemented with 10% fetal bovine serum (FBS) (ThermoFisher) and 1% penicillin-streptomycin (ThermoFisher) at 37 °C with 5% CO_2_. The WT SARS-CoV-2 virus (GDPCC-nCOV01, GISAID: EPI_ISL_403934), the B.1.351 variant (SARS-CoV-2/human/CHN/20SF18530/2020 strain; GDPCC-nCOV84, National Genomics Data Center: ACC_GD530_GZ_2020-12-10) and the B.1.617.2 variant (GDPCC 2.00096, sequence has not been published) were obtained from the Guangdong Center for Human Pathogen Culture Collection (GDPCC) at Guangdong Provincial Center for Disease Control and Prevention. The D614G variant (hCoV-19/CHN/SYSU-IHV/2020 strain; GISAID: EPI_ISL_444969) was isolated from the sputum of a female infected individual and propagated in Vero E6 cells.

### Surface Plasmon Resonance (SPR) assay

SPR experiments were performed using a Biacore® T100 instrument (GE Healthcare, Uppsala, Sweden). All binding analyses were performed at 25 °C using HBS-EP+ (10 mM HEPES, 150 mM NaCl, 3 mM EDTA, 0.005% Tween-20) as the running buffer. Experiments were executed following the previously described SPR protocol^45^. Briefly, RBD protein was immobilized on the sensor chip CM5-type surface raising a final immobilization level of ∼600 Resonance Units (RU). Serial dilutions of 35B5 were injected in concentration from 10 to 0.625 nM. Serial dilutions of 32C7 were injected in concentration from 120 to 7.5 nM. For the competitive binding assays, the first sample flew over the chip at a rate of 20 µl/min for 120 s, then the second sample was injected at the same rate for another 200 s. The response units were recorded at room temperature and analyzed using the same software as mentioned above.

### ELISA

The ELISA plates were coated with 50 ng of SARS-CoV-2 RBD protein (Sino Biological, 40592-V08H) in 100 µl PBS per well overnight at 4 °C. On the next day, the plates were incubated with blocking buffer (5% FBS + 0.1% Tween 20 in PBS) for 1 hour. Then, serially diluted mAbs in 100 µl blocking buffer were added to each well and incubated for 1 hour. After washing with PBST (PBS + 0.1% Tween 20), the bound antibodies were incubated with HRP-conjugated goat anti-human IgG antibody (Bioss Biotech) for 30 min, followed by washing with PBST and addition of TMB (Beyotime). The ELISA plates were allowed to react for ∼5 min and then stopped by 1 M H_2_SO_4_ stop buffer. The optical density (OD) value was determined at 450 nm. EC_50_ values were determined by using Prism 6.0 (GraphPad) software after log transformation of the mAb concentration using sigmoidal dose-response nonlinear regression analysis.

### Neutralization assay

Infectious SARS-CoV-2 neutralization assay was performed according to previous reports^46^. Vero E6 cells were seeded in a 24-well culture plates at a concentration of 4×10^4^ cells per well at 37 °C for 24 h. For infection with authentic SARS-CoV-2 at an MOI of 0.005, 200 µl of diluted authentic SARS-CoV-2 and 5-fold serially diluted 35B5 or 32C7 (from 5 µg/mL to 0.064 µg/mL) mAbs were mixed in the medium with 2% FBS, and were then added into the Vero E6 cells. The culture supernatant was collected at 48h post-infection for focus forming assay and qRT-PCR. IC_50_ values were determined by nonlinear regression using Prism 6.0 (GraphPad).

### Focus forming assay (FFA)

The virus titer was detected by FFA, which is characterized by its high-throughput as compared to the traditional plaque assay^47^. Briefly, Vero E6 cells were seeded in 96-well plates 24h prior to infection. Virus cultures were serially diluted and used to inoculate Vero E6 cells at 37°C for 1 h, followed by changed with fresh medium containing 1.6% carboxymethylcellulose. After 24 hours, Vero E6 cells were fixed with 4% paraformaldehyde and permeabilized with 0.5% Triton X-100. Cells were then incubated with anti-SARS-CoV-2 nucleocapsid protein polyclonal antibody (Sino Biological), followed by an HRP-labeled secondary antibody (Proteintech). The foci were visualized by TrueBlue Peroxidase Substrate (SeraCare Life Science), and counted with an ELISPOT reader.

### Protection against SARS-CoV-2 in hACE2 mice

All animal experiments were carried out in strict accordance with the guidelines and regulations of Laboratory Monitoring Committee of Guangdong Province of China and were approved by Ethics Committee of Zhongshan School of Medicine of Sun Yat-sen University on Laboratory Animal Care (SYSU-IACUC-2021-00432). Viral infections were performed in a biosafety level 3 (BSL3) facility in accordance with recommendations for the care and use of laboratory animals. The hACE2 mice of the same sex were randomly assigned to each group. For infection, ICR-hACE2 mice were anesthetized with isoflurane and inoculated intranasally with 4×10^4^ PFU SARS-CoV-2 virus (GISAID: EPI_ISL_402125). Six hours later, the infected mice received a single dose of 35B5 (20 mg/kg) or 32C7 (20 mg/kg) or vehicle. And the H11-K18-hACE2 mice were intranasally challenged with 4×10^4^ PFU of three subtypes SARS-CoV-2 virus (D614G, B.1.351 and B.1.617.2) per mouse, respectively. Four hours later, the infected mice received a single dose of 35B5 (30 mg/kg) or vehicle. The lungs were collected at day 5 post-infection for further assays.

### Measurement of viral burden

For *in vitro* neutralization assay, RNA of culture supernatant was extracted by using TRIzol reagent (Invitrogen). For *in vivo* neutralization assay, lungs of SARS-CoV-2 infected mice were collected and homogenized with gentle MACS M tubes (Miltenyi Biotec, 130-093-236) in a gentle MACS dissociator (Miltenyi Biotec, 130-093-235). Then, total RNA of homogenized lung tissues was extracted with RNeasy Mini Kit (QIAGEN, 74104) according to the manufacturer’s instruction. The extracted RNA was performed with quantitative RT-PCR (qRT-PCR) assay to determine the viral RNA copies by using one-step SARS-CoV-2 RNA detection kit (PCR-Fluorescence Probing) (Da An Gene Co., DA0931).

To generate a standard curve, the SARS-CoV-2 nucleocapsid (N) gene was cloned into a pcDNA3.1 expression plasmid and *in vitro* transcribed to obtain RNAs for standards. Indicated copies of N standards were 10-fold serially diluted and proceeded to qRT-PCR utilizing the same one-step SARS-CoV-2 RNA detection kit to obtain standard curves. The reactions were carried out on a QuantStudio 7 Flex System (Applied Biosystems) according to the manufacturer’s instruction under the following reaction conditions: 50 °C for 15 min, 95 °C for 15 min, and 45 cycles of 94 °C for 15 s and 55 °C for 45 s. The viral RNA copies of each tissue were calculated into copies per ml and presented as log10 scale. The N specific primers and probes were: N-F (5’- CAGTAGGGGAACTTCTCCTGCT-3’), N-R (5’-CTTTGCTGCTGCTTGACAGA-3’) and N-P (5’-FAM-CTGGCAATGGCGGTGATGCTGC-BHQ1-3’). In each qRT-PCR experiment, both positive control and negative control of simulated RNA virus particles were included to monitor the entire experimental process and ensure the reliability of the test results.

### Quantification of cytokine and chemokine mRNA

RNA was isolated from lung homogenates as described above. Then, cDNA was synthesized from isolated RNA using HiScript III RT SuperMix for qPCR (Vazyme Biotech). The mRNA expression levels of cytokine and chemokine were determined by using ChamQ Universal SYBR q-PCR Master Mix (Vazyme Biotech) with primers for *IL-6* (forward: CCCCAATTTCCAATGCTCTCC; reverse: CGCACTAGGTTTGCCGAGTA), *IL-1b* (forward: TCGCTCAGGGTCACAAGAA; reverse: GTGCTGCCTAATGTCCCCTT), *Tnfa* (forward: ATGGCTCAGGGTCCAACTCT; reverse: CGAGGCTCCAGTGAATTCGG), Ccl2 (forward: AACTGCATCTGCCCTAAGGT; reverse: AGGCATCACAGTCCGAGTCA) and *Cxcl1* (forward: ACTCAAGAATGGTCGCGAGG; reverse: GTGCCATCAGAGCAGTCTGT). The results were normalized to *Gapdh* levels.

### Histopathology

At day 5 post-SARS-CoV-2 infection, hACE2 mice were euthanized, and lungs were collected and fixed in 4% paraformaldehyde buffer. Tissues were embedded with paraffin and sections (3-4 mm) were stained with hematoxylin and eosin (H&E).

### Production and purification of S-2P and S-6P proteins

The plasmids encoding the ectodomains of the SARS-CoV-2 S-2P and S-6P mutants were kindly provided by Dr. Junyu Xiao. HEK293F cells were cultured in the SMM 293-TI medium (Sino Biological Inc.) at 37 °C with 8% CO_2_. The S-2P and S-6P plasmids were transiently transfected into HEK293F cells using 25-kDa linear polyethyleneimine (PEI) (Polysciences) with the PEI:DNA mass ratio of 3:1 and 1 mg DNA for per liter of culture when the cell density reached 2 × 10^6^ cells per mL. At day 4 post-transfection, the supernatants of the cell culture were harvested by centrifugation at 10,000 × g for 30 min. The secreted S-2P and S-6P proteins were purified using HisPur^TM^ cobalt resins (Thermo Scientific) and StrepTactin resins (IBA). Further purification was carried out using size-exclusion chromatography with a Superose 6 10/300 column (GE Healthcare) in the buffer containing 20 mM HEPES pH 7.2, 150 mM NaCl and 10% Trehalose. The Fab regions of the 35B5 and 32C7 were obtained after the digestion by papain for 40 min at 37 °C in a buffer containing 20 mM HEPES pH 7.2, 150 mM NaCl, 5 mM EDTA and 5 mM L-cysteine. The obtained Fabs were purified with a Desalting column (GE Healthcare Life Sciences) to remove L-cysteine, and then further purified in a HiTrap Q column (GE Healthcare Life Sciences). The purified Fabs were collected and concentrated to 0.6 mg/mL.

### Negative staining analysis

For negative-staining assays, the S-2P, S-6P, 35B5 Fab and 32C7 Fab proteins were diluted to 0.02 mg/ml in the buffer of 20 mM HEPES, pH 7.2, and 150 mM NaCl. 2 µL of 35B5 Fab or 32C7 Fab was mixed with 2 µL S-2P or S-6P and incubated on ice for 3 and 10 min, respectively, or at room temperature for 30 min. The samples were loaded in the glow-discharged carbon-coated copper grids and stained with 3% Uranyl Acetate (UA). The prepared grids were examined using a Tecnai G2 Spirit BioTWIN transmission electron microscope (FEI) operated at 120 kV. Micrographs were recorded and analyzed using Digital Micrograph software with 120,000× nominal magnification.

### Cryo-EM sample preparation and data collection

2 μL of S-6P (1.2 mg/mL) and 2 μL of 35B5 Fab or 32C7 Fab (0.6 mg/mL) were incubated for 3 min at room temperature and then loaded onto the glow-discharged holy-carbon gold grids (Quantifoil, R1.2/1.3). The grids were washed using the buffer containing 20 mM HEPES, pH 7.2, and 150 mM NaCl. The washed grids were blotted using a Mark IV Vitrobot (Thermo Fisher) at 100% humidity and 16 °C for 3 s, and *submerged in liquid ethane by plunge-freezing.* For the S-6P-35B5 Fab complex, micrographs were recorded on a FEI Titan Krios (Thermo Fisher) electron microscope operated at 300 kV. Totally, 3740 movies were recorded on a K3 Summit direct electron detector (Gatan) in the super-resolution mode (0.5475 Å/pixel) at a nominal magnification of 81,000 using a defocus range of 1.2 to 1.3 μm. A GIF Quantum energy filter (Gatan) with a slit width of 20 eV was used on the detector. The micrographs were dose-fractioned into 32 frames with a total electron exposure of ∼50 electrons per Å^2^.

For the S-6P-32C7 Fab complex, micrographs were collected on a FEI Titan Krios (Thermo Fisher) operating at 300 kV using the AutoEMation software ^48^. Totally, 2528 movies were recorded on a K3 Summit direct electron detector (Gatan) in the super-resolution mode (0.53865 Å/pixel) at a nominal magnification of 81,000 using a defocus range of 1.4 to 1.8 µm. A GIF Quantum energy filter (Gatan) with a slit width of 20 eV was used on the detector. The micrographs were dose-fractioned into 32 frames with a total electron exposure of ∼50 electrons per Å^2^.

### Cryo-EM image processing

Raw movie frames were binned, aligned and averaged into motion-corrected summed images using MotionCor2^49^. The dose-weighted images were then imported into cryoSPARC^50^ for the following image processing, including CTF estimation, particle picking and extraction, 2D classification, *ab initio* 3D reconstruction, heterogeneous 3D refinement and non-uniform homogeneous refinement. For the S-6P-35B5 Fab complex, 8 representative particle templates were generated in 2D classification of 65,242 particles auto-picked by the blob picker from the first 1000 micrographs. Using these templates, 1,178,527 particles were extracted with a box size of 330×330 and classified into 150 classes in 2D classification.

Among them, 43 classes that included 818,470 particles were selected for *ab initio* 3D reconstruction and heterogeneous refinement. Finally, 392,378 particles reconstructed an apparent architecture of the S-6p-35B5 Fab complex and were subjected to two more rounds of *ab initio* 3D reconstruction and heterogeneous refinement before non-uniform refinement. Then the particles were subjected to global and local CTF refinement for final non-uniform refinement, and generated three abundant populations of the S-6P-35B5 Fab complex and structures. Local refinements of the RBD-35B5 Fab region were performed to improve the interface density in *cryoSPARC*. Sharpened maps were generated and validated for model building and refinement. Reported resolutions are based on the gold-standard Fourier shell correlation^51, 52^ of 0.143 criterion.

For the S-6P-32C7 Fab complex, 4 representative particle templates were generated in 2D classification of 122,851 particles auto-picked by the blob picker from the first 1000 micrographs. Based on these templates, 744,865 particles were extracted with a box size of 380×380 and classified into 150 2D classes. Among them, 28 classes including 338,454 particles were selected for *ab initio* 3D reconstruction and heterogeneous refinement. 125,858 particles that reconstructed an apparent architecture of the S-6P-32C7 Fab complex were subjected to two more rounds of *ab initio* 3D reconstruction and heterogeneous refinement before non-uniform refinement. Finally, the S-6P-32C7 Fab complex structure that only includes a bound 32C7 Fab was reconstructed from 119,062 particles. Local refinement of the RBD-32C7 Fab region was also performed.

### Cryo-EM structure modeling and analysis

To build the S-6P-35B5 Fab complex structural model, an “up” RBD-35B5 Fab model were first generated using a Fab structure (PDB ID: 2X7L) and a RBD model from the Spike trimer (PDB ID: 7K8Y) and manually built in Coot^53^ in the locally refined map. The obtained RBD-35B5 Fab model was superimposed with the intact Spike trimer structure (PDB ID: 7K8Y) to generate an initial model of the S-6P-35B5 Fab complex model. The “down” RBD-35B5 Fab model was obtained from fitting the structures of 35B5 Fab, the RBD and NTD domains into the locally refined map of the the S-6P-35B5 Fab complex of State 1 and then validated by *Phenix*^54^. The model building of the S-6P-32C7 Fab complex was carried out in the similar procedure as that of the S-6P-35B5 Fab complex with a Spike trimer structure (PDB ID: 6XKL) as a template. All model building were performed in Coot^53^. Structural refinement and validation were carried out in *Phenix*^54^. Structural figures were generated using UCSF ChimeraX version 1.2^55^.

### Statistics

In the mouse study assessing mAb protection against WT SARS-CoV-2, the comparisons of lung viral titers and lung cytokine/chemokine mRNA were performed using one-way ANOVA with Tukey’s post hoc test by Prism 6.0 (GraphPad). In the mouse study assessing mAb protection against SARS-CoV-2 VOCs, the comparison of lung viral titers was performed using t test (unpaired) by Prism 6.0 (GraphPad).

### Data and materials availability

The cryo-EM maps and atomic coordinates have been deposited to the Electron Microscopy Data Bank (EMDB) and Protein Data Bank (PDB) with accession codes EMD-31033 and PDB 7E9N (State 1 of the 35B5 Fab-S-6P complex), EMD-31444 and PDB 7F46 (“down” RBD/35B5 Fab local refinement of State 1), EMD-31034 and PDB 7E9O (State 2 of the 35B5 Fab-S-6P complex), EMD-31035 and PDB 7E9P (“up” RBD/35B5 Fab local refinement of State 2), EMD-31036 and PDB 7E9Q (State 3 of the 35B5 Fab-S-6P complex), EMD-31209 and PDB 7ENF (the 32C7 Fab-S-6P complex), and EMD-31210 and PDB 7ENG (“down” RBD/32C7 Fab local refinement).

## Acknowledgements

We thank Guangdong Center for Human Pathogen Culture Collection (GDPCC) for providing SARS-CoV-2 isolates. We thank Dr. Junyu Xiao (Peking University) for providing the plasmids encoding the ectodomains of the SARS-CoV-2 S-2P and S-6P mutants. This work was supported by grants from the National Natural Science Fund for Distinguished Young Scholars (No. 31825011 to L.Y.), the National Science and Technology Major Project (No. 2017ZX10202102-006-002 to L.Y.), Guangdong Innovative and Entrepreneurial Research Team Program (2016ZT06S638 to K.D.), High-level Biosafety Laboratory Construction and Operation Program of the Science and Technology Projects of Guangdong Province of China to K.D., the National Natural Science Fund (81925024 to Y. Z.), the National Key Research and Development Program of China (2017YFA0503900 to Y. Z.), and the Fundamental Research Funds for the Central Universities to Y. Z..

## Author contributions

X.C., S.Y., Z.P., Y.Y., Y.L., J.Z., L.G., J.Z., L.X. and Q.H. collected the PBMC, isolated SARS-CoV-2 RBD-specific B cells and cloned the antibodies; A.H., Y.Z., F.Y., J.Z., F.L., Y.S., F.H., X.Y., Y.P., L.T., H.Z., H.Z., J.H. and H.Z. performed *in vitro* and *in vivo* SARS-CoV-2 virus neutralization assays; X.W. and A.L. performed negative stainings and cryo-EM analyses. L.Y. designed the study and analyzed the data with Y.Z., K.D., X.C., X.W. and A.H.; X.C. wrote the first draft of the manuscript, which was edited by Y.Z., L.Y., K.D., X.W. and A.H.; Y.Z., K.D. and L.Y. supervised the study.

## Competing interests

The pending patents of 35B5 and 32C7 have been licensed.

---

**Supplementary Figure 1.**
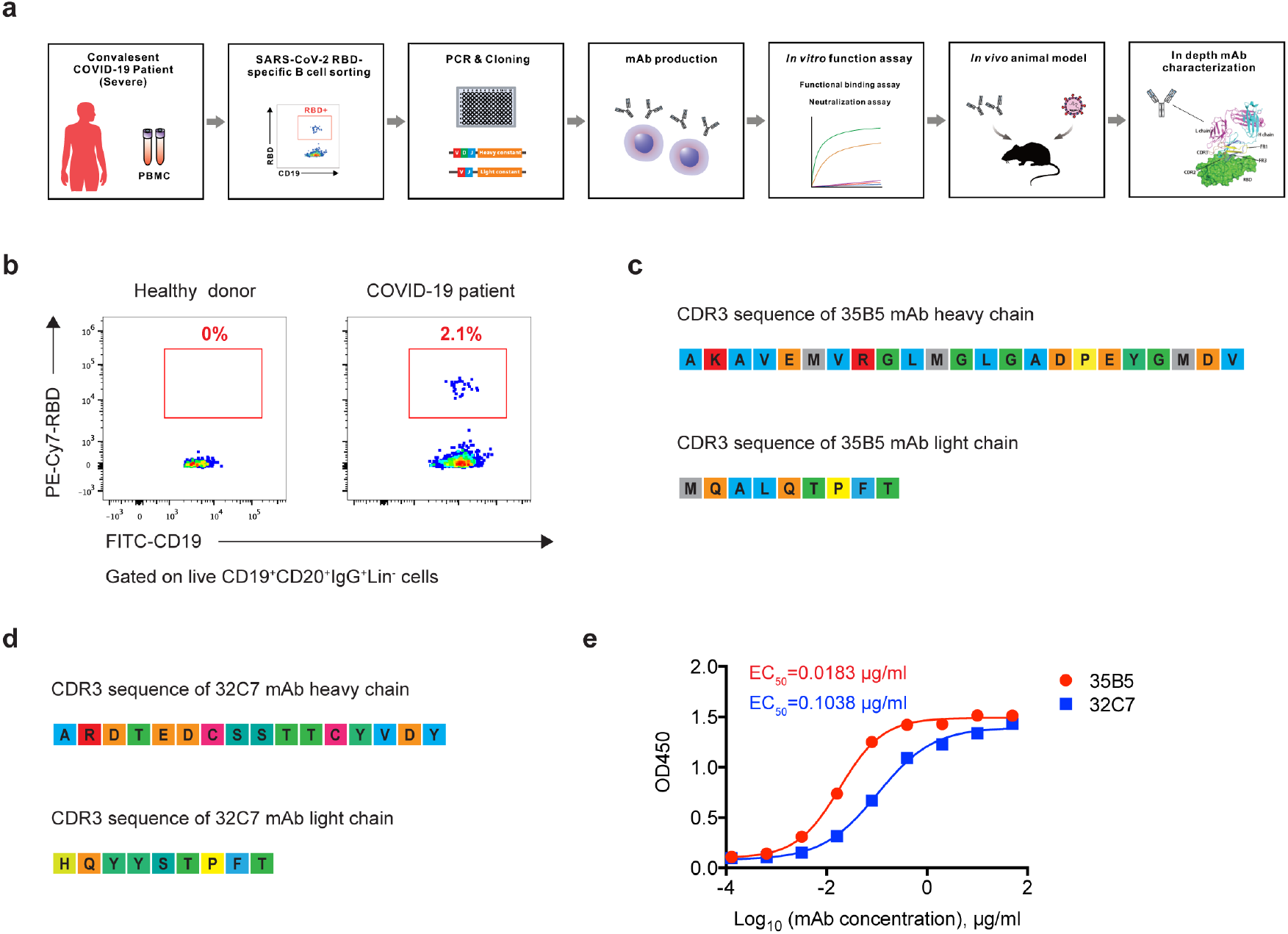
Isolation of potent SARS-CoV-2 neutralizing mAbs from COVID-19 convalescent patients. **a** Isolation strategy of SARS-CoV-2 neutralizing mAbs. **b** Flow cytometry analysis of SARS-CoV-2 RBD-specific B cells from the PBMC of healthy donors and COVID-19 convalescent patients. The numbers adjacent to the outlined area indicate the proportions of SARS-CoV-2 RBD-specific B cells in CD19^+^CD20^+^IgG^+^ B cells. **c, d** The CDR3 sequences of mAb 35B5 (**c**) and mAb 32C7 (**d**). **e** ELISA analysis of mAb 35B5 (red) or mAb 32C7 (blue) binding to SARS-CoV-2 RBD protein. EC_50_, concentration for 50% of maximal effect.

**Supplementary Fig. 2.**
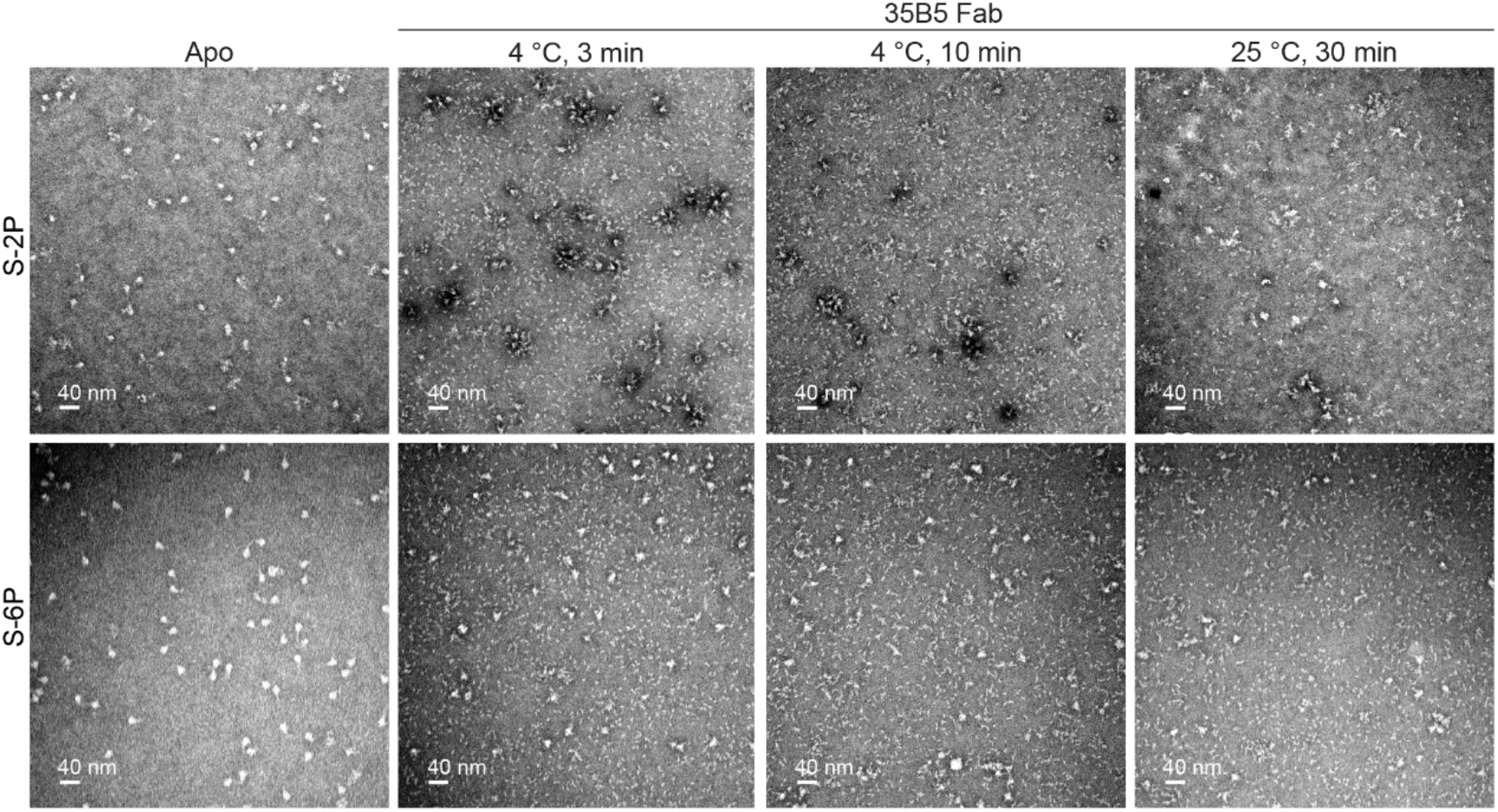
Representative negative-staining EM micrographs of the S-2P and S-6P trimers after the treatment of 35B5 Fab. The S-2P and S-6P trimeric proteins were treated with or without 35B5 Fab for indicated time at 4°C or 25°C before negative staining analysis. Scale bar, 40 nm.

**Supplementary Fig. 3.**
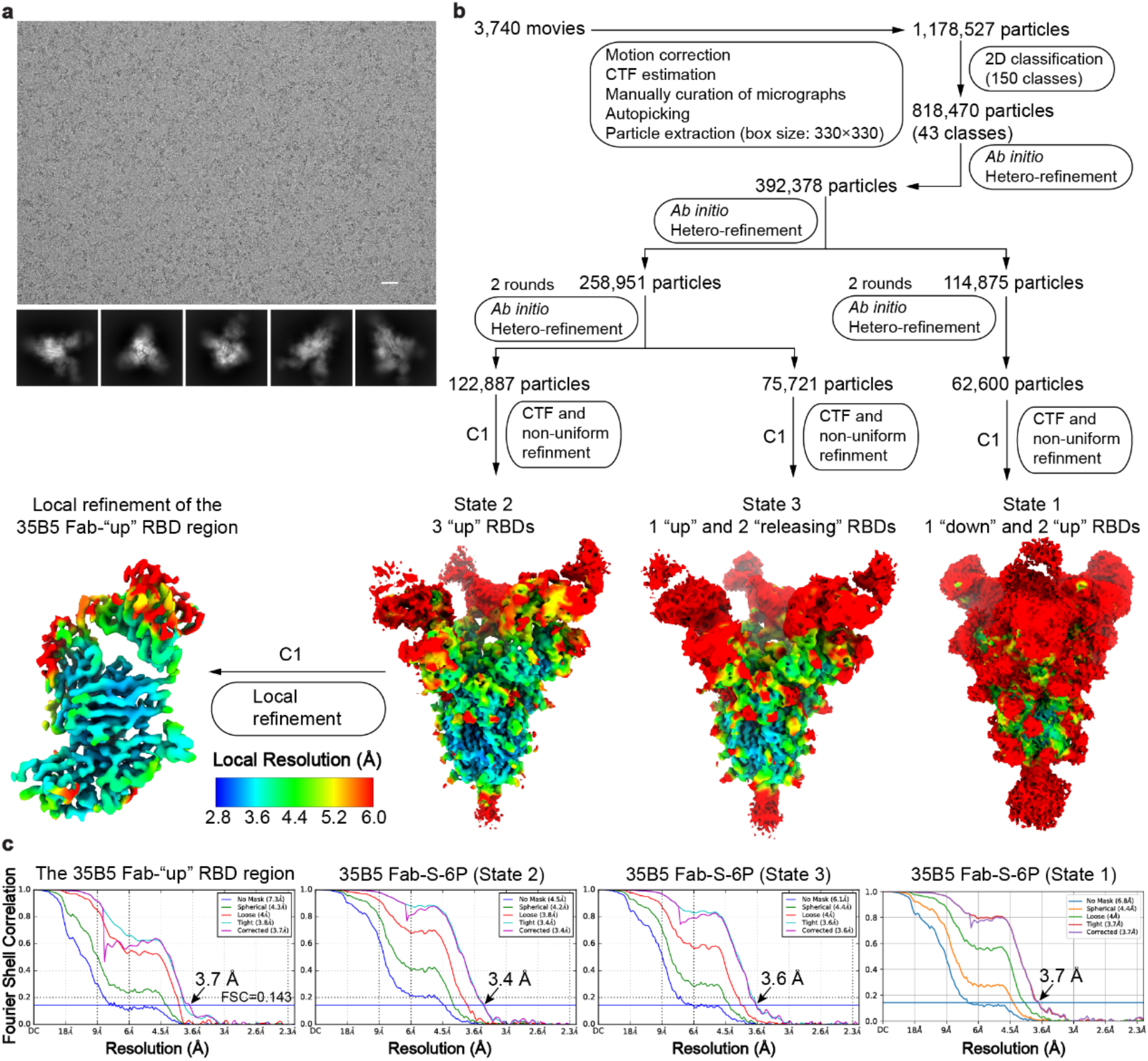
Cryo-EM data processing and validation of the S-6P-35B5 Fab complex. **a** Representative cryo-electron micrograph (upper) and 2D class averages (lower) of the S-6P-35B5 Fab complex. Scale bar, 25 nm. **b** Cryo-EM data processing flow-chart of the S-6P-35B5 Fab complex. Three states of the S-6P-35B5 Fab complex were obtained in data collection. Local refinement of the 35B5Fab-“up” RBD region was carried out using C1 symmetry (left). All density maps were prepared using ChimeraX. The map resolution is color coded for different regions. The resolution goes from 2.8 to 6.0 Å. **c** The Fourier shell correlation (FSC) curves for the reconstructions in **b**. The resolution estimations of cryo-EM density maps were based on the corrected FSC curves at the gold standard FSC=0.143 criterion.

**Supplementary Fig. 4.**
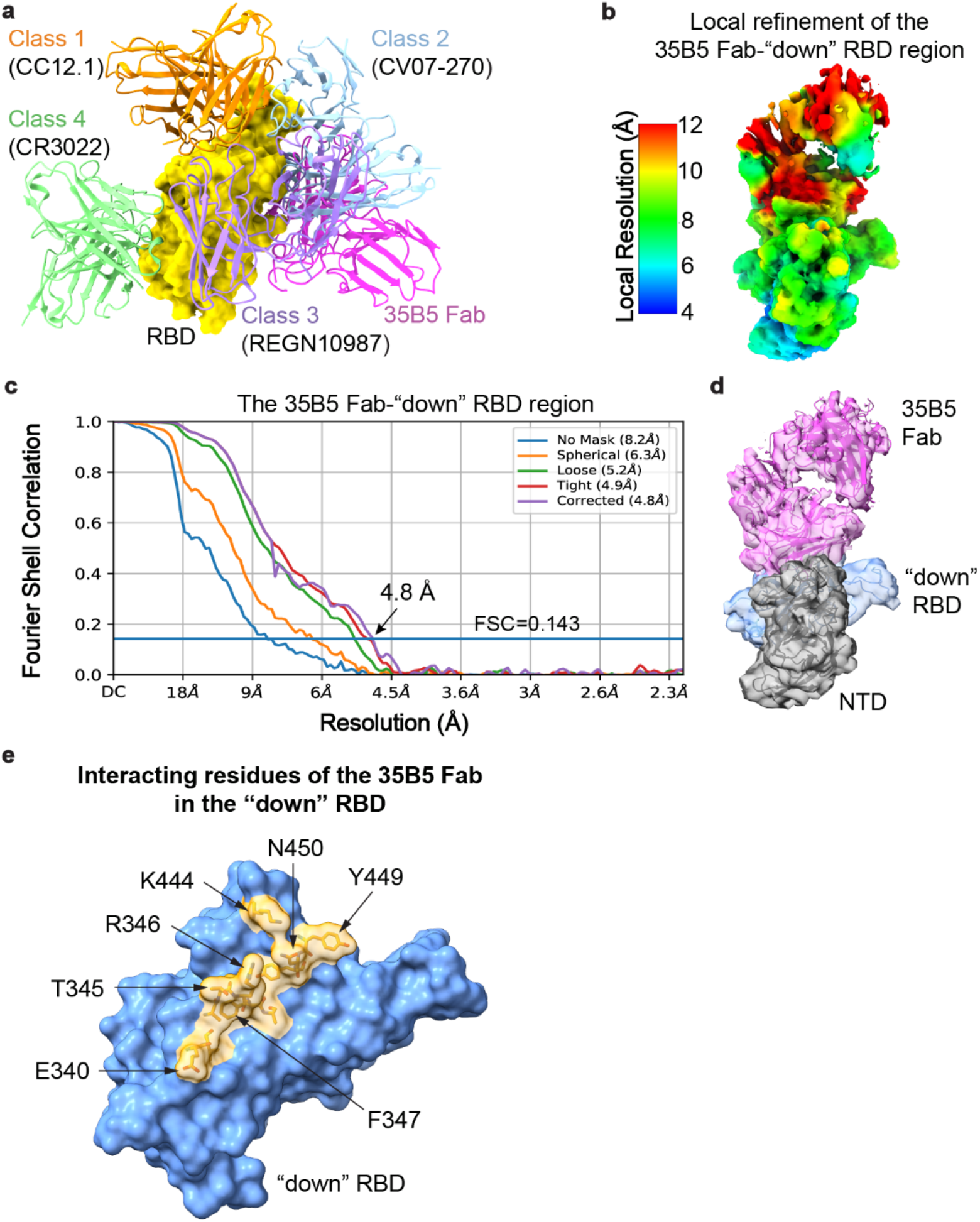
Comparison of 35B5 with other mAbs and the local refinement of 35B5 Fab-“down” RBD region. **a** Structural comparisons of the 35B5 Fab-“up” RBD interactions with those of the previously identified four classes of neutralizing antibodies. 35B5 Fab and the Fab regions of the representative antibodies CC12.1, CV07-270, REGN10987 and CR3022 from Class 1-4, respectively, are illustrated in cartoon and colored as indicated. RBD is represented as surface in yellow. **b** Local refinement of the region of 35B5 Fab with the “down” RBD and NTD domains in the State 1 S-6P-35B5 Fab complex. The density map was prepared using ChimeraX. The map resolution is color coded for different regions. The resolution goes from 4.0 Å to 12 Å. **c** The Fourier shell correlation (FSC) curve for the reconstructions in **b**. The resolution estimation (4.8 Å) of cryo-EM density maps were based on the corrected FSC curves at the gold standard FSC=0.143 criterion. **d** Structural modeling of 35B5 Fab with the “down” RBD and NTD domains in the density map obtained from the local refinement in **b**. 35B5 Fab, RBD and NTD are colored in purple, blue and grey respectively. **e** The 35B5-interacting residues of the “down” RBD. The 35B5-interacting residues of the “down” RBD (within 4 Å to 35B5 Fab) are colored in yellow.

**Supplementary Fig. 5.**
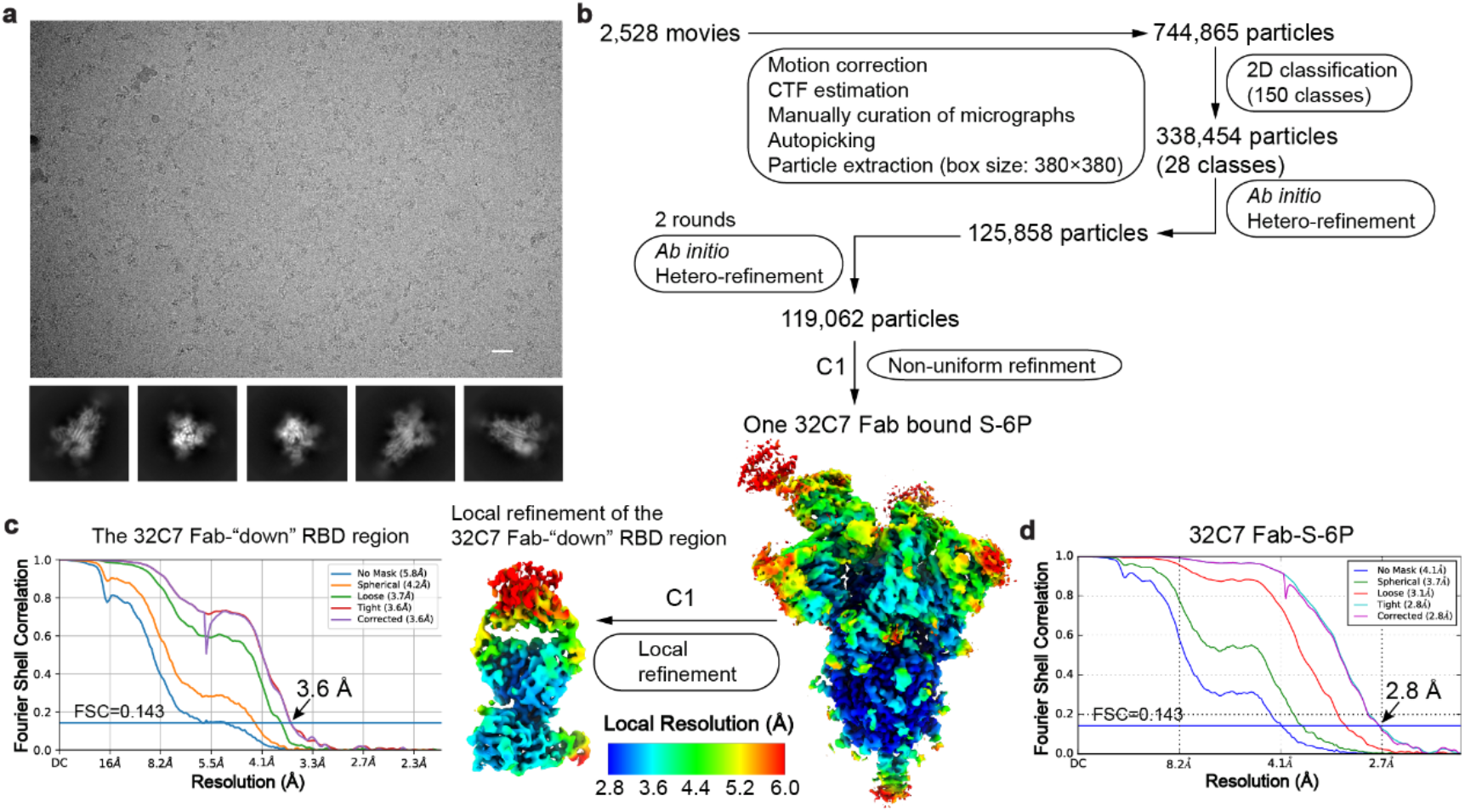
Cryo-EM data processing and validation of the S-6P-32C7 Fab complex. **a** Representative electron micrograph (upper) and 2D class averages (lower) of the SARS-CoV-2 S-6P- 32C7 Fab complex. Scale bar, 25 nm. **b** The cryo-EM data processing flow chart of the S-6P-32C7 Fab complex. Only one 32C7 Fab is bound to a “down” RBD of the S-6P trimer. Local refinement of 32C7 Fab with the “down” RBD generated a map at the resolution of ∼ 3.6 Å. The density map was prepared using ChimeraX. The map resolution is color coded for different regions. The resolution goes from 2.8 Å to 6.0 Å. **c**, **d** The FSC curves for the reconstructions of the 32C7 Fab-RBD region (**c**) and the S-6P-32C7 Fab complex (**d**) in **b**. The resolution estimations of cryo-EM density maps were based on the corrected FSC curves at the gold standard FSC=0.143 criterion.

**Supplementary Fig. 6.**
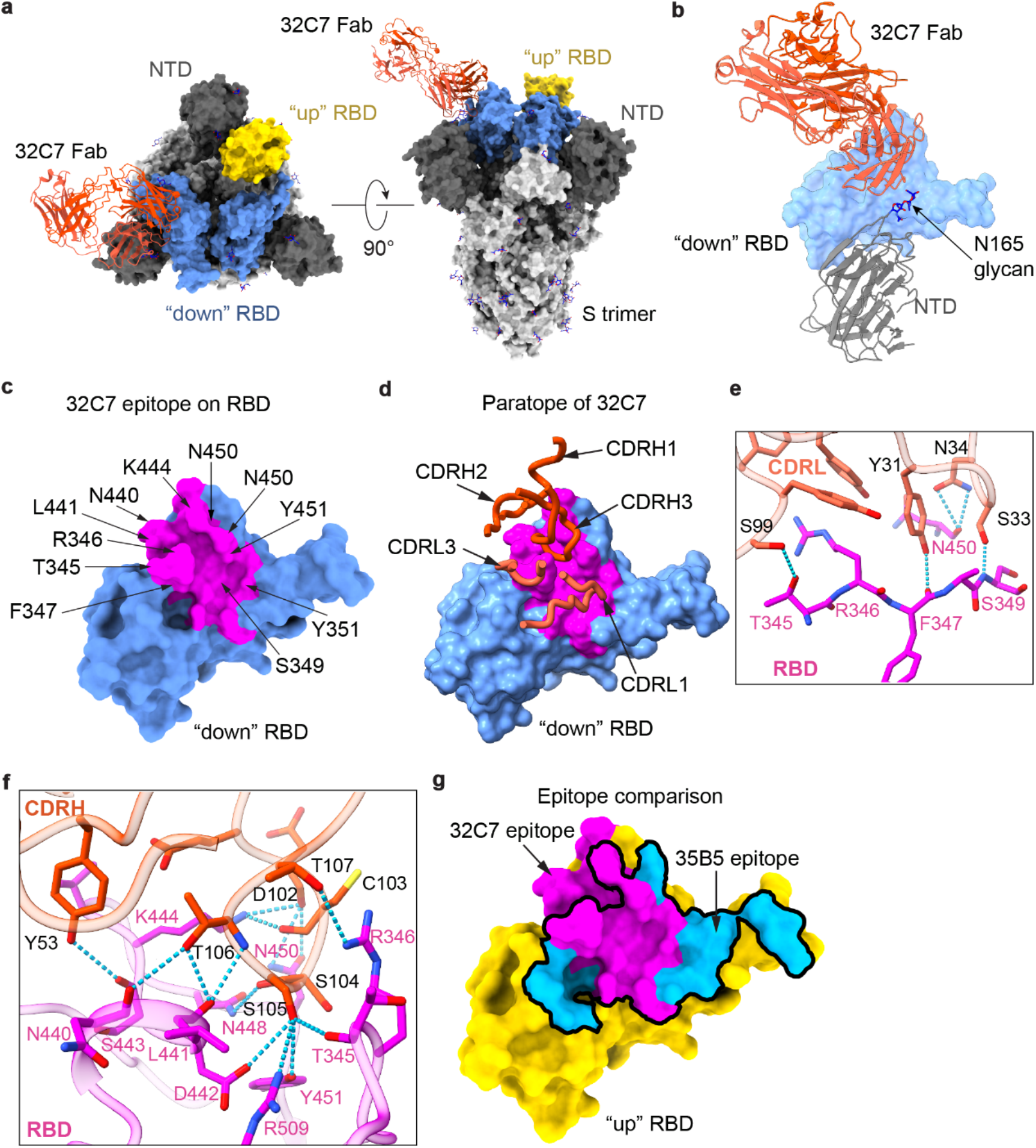
Cryo-EM structure of the S-6P-32C7 Fab complex. **a** The 2.8-Å cryo-EM structure of the S-6P-32C7 Fab complex. The S-6P trimer is represented as surface. 32C7 Fab is shown in cartoon in red. The “down” and “up” RBDs are colored in blue and yellow, respectively. **b** Structural superposition of the 32C7 Fab-RBD model and the tight-closed S trimer (PDBID: 6ZB5). In the structural superimposition, 32C7 Fab does not form structural clashes with the NTD domain of the tight-closed S trimer. RBD is shown as surface in blue. The NTD domain of the S trimer is shown in cartoon in grey. **c** The 32C7 epitope on RBD. The epitope residues are labeled as indicated. **d** The RBD-interacting regions in 32C7 Fab. The CDRH1, CDRH2, CDRH3, CDRL1 and CDRL3 are shown in ribbon. **e**, **f** Detailed interactions of the light chain CDRs (**e**) and heavy chain CDRs (**f**) of 32C7 Fab with RBD. Hydrogen-bond interactions are shown as dashed lines. **g** Comparison of the epitopes for 32C7 and 35B5 on the surface of RBD. The epitopes for 32C7 and 35B5 (black contour) are colored in purple and blue, respectively.

**Supplementary Fig. 7.**
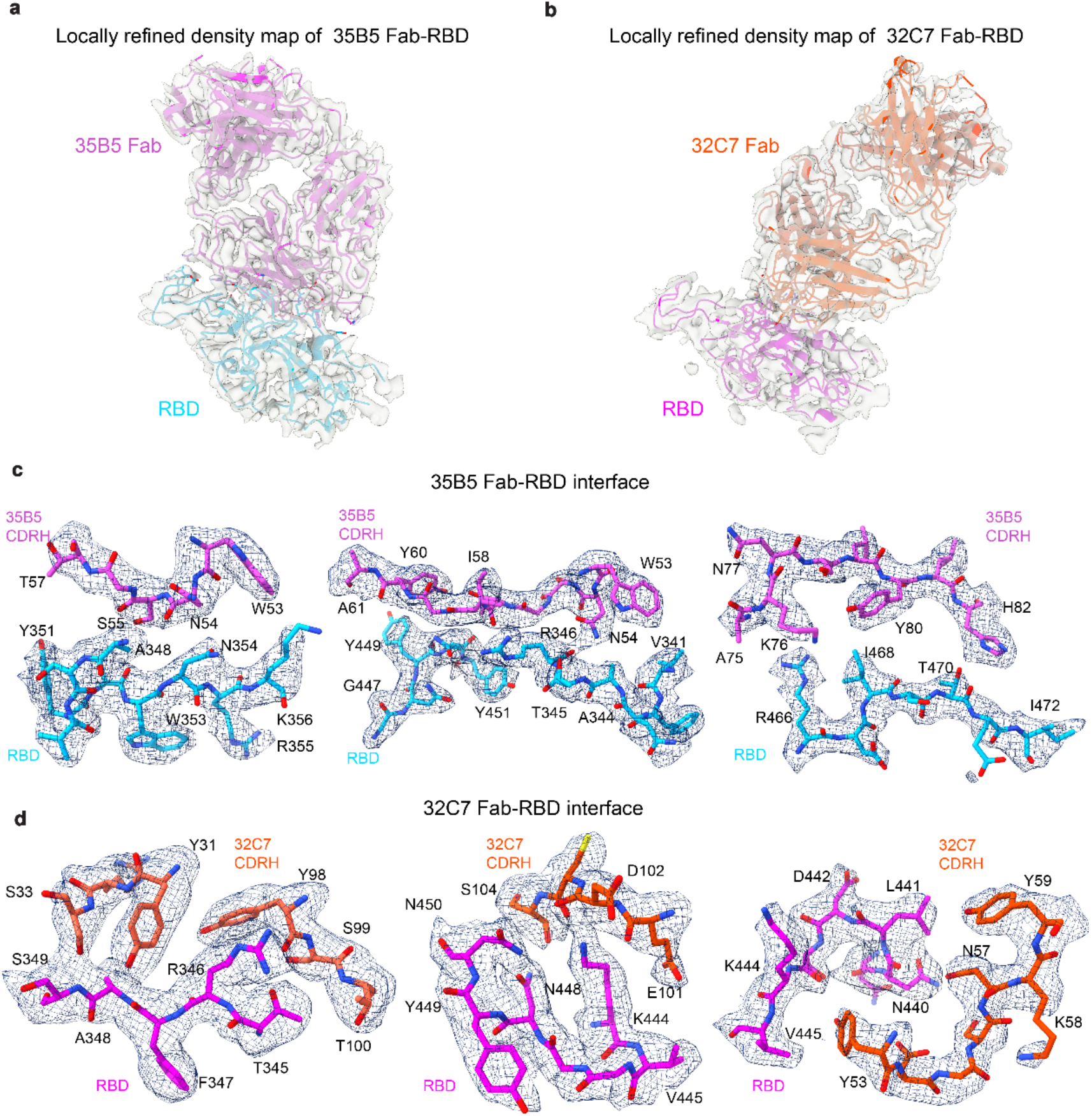
Density maps of 35B5 Fab-RBD and 32C7 Fab-RBD interfaces after local refinement. **a** Locally refined density map (transparent gray) of the 35B5 Fab-“up” RBD region in the State 2 S-6P-35B5 Fab complex. **b** Locally refined density map of the 32C7 Fab-RBD region in the S-6P-32C7 Fab complex. **c** Representative density maps of interaction regions at the 35B5 Fab-RBD interface. **d** Representative density maps of interaction regions at 32C7 Fab-RBD interface. The interacting residues of RBD and Fabs are shown as stick.

**Supplementary Table 1.** Cryo-EM data collection and refinement statistics.

|  | 35B5 Fab-S-6P<br>(state 1) | 35B5 Fab-S-6P<br>(state 1, local<br>refinement) | 35B5 Fab-S-6P<br>(state 2) | 35B5 Fab-S-6P<br>(state 2, local<br>refinement) | 35B5 Fab-S-6P<br>(state 3) | 32C7 Fab-S-6P | 32C7 Fab-S-6P<br>(local<br>refinement) |
| --- | --- | --- | --- | --- | --- | --- | --- |
| **PDB code** | **7E9N** | **7F46** | **7E9O** | **7E9P** | **7E9Q** | **7ENF** | **7ENG** |
| **EMDB code** | **EMD-31033** | **EMD-31444** | **EMD-31034** | **EMD-31035** | **EMD-31036** | **EMD-31209** | **EMD-31210** |
| **Data collection and processing** |  |  |  |  |  |  |  |
| Voltage (kV) | 300 | 300 | 300 | 300 | 300 | 300 | 300 |
| Magnification | 81,000 | 81,000 | 81,000 | 81,000 | 81,000 | 81,000 | 81,000 |
| Pixel size (Å/pix) | 0.5475 | 0.5475 | 0.5475 | 0.5475 | 0.5475 | 0.53865 | 0.53865 |
| Frames per exposure | 32 | 32 | 32 | 32 | 32 | 32 | 32 |
| Exposure (e<sup>-</sup>/Å<sup>2</sup>) | 50 | 50 | 50 | 50 | 50 | 50 | 50 |
| Defocus range (µm) | 1.2 to 1.3 | 1.2 to 1.3 | 1.2 to 1.3 | 1.2 to 1.3 | 1.2 to 1.3 | 1.4 to 1.8 | 1.4 to 1.8 |
| Final particle images (no.) | 62,600 | 62,600 | 122,887 | 122,886 | 75,721 | 119,062 | 119,062 |
| Symmetry imposed | C1 | C1 | C1 | C1 | C1 | C1 | C1 |
| Map resolution (Å, 0.143 FSC threshold) | 3.7 | 4.8 | 3.4 | 3.7 | 3.6 | 2.8 | 3.6 |
| **Refinement** |  |  |  |  |  |  |  |
| Map resolution (Å, 0.5 FSC threshold) | 3.9 | 8.6 | 3.7 | 3.9 | 3.8 | 3.0 | 3.7 |
| Map sharpening *B* factor (Å<sup>2</sup>) | 81.9 | 115.6 | 90.4 | 86.3 | 82.5 | 75.8 | 110.2 |
| **Model composition** |  |  |  |  |  |  |  |
| Protein residues | 4,313 | 842 | 4338 | 639 | 4,338 | 3,390 | 640 |
| Ligands | 27 | 2 | 27 | 1 | 27 | 53 | 2 |
| **R.m.s. deviations** |  |  |  |  |  |  |  |
| Bond length (Å) | 0.008 | 0.005 | 0.004 | 0.005 | 0.005 | 0.005 | 0.006 |
| Bond angles (°) | 0.870 | 0.757 | 0.768 | 0.849 | 0.887 | 0.835 | 0.802 |
| **Ramachandran plot** |  |  |  |  |  |  |  |
| Favored (%) | 91.61 | 91.12 | 95.12 | 92.53 | 93.60 | 94.70 | 92.59 |
| Allowed (%) | 8.39 | 8.88 | 4.88 | 7.47 | 6.40 | 5.30 | 7.41 |
| Outliers (%) | 0 | 0 | 0 | 0 | 0 | 0 | 0 |
| **Validation** |  |  |  |  |  |  |  |
| Poor rotamers (%) | 0.11 | 0 | 0.24 | 0 | 0.32 | 0.56 | 0.36 |
| Clash score | 12.95 | 18.16 | 10.62 | 11.97 | 14.62 | 10.14 | 14.32 |
| MolProbity score | 2.12 | 2.27 | 1.88 | 2.06 | 2.09 | 1.89 | 2.13 |

## References

1. World Health Organization. Weekly epidemiological update on COVID-19 - 26 October 2021. https://www.who.int/publications/m/item/weekly-epidemiological-update-on-covid-19---26-october-2021.

2. Corti D, Purcell LA, Snell G, Veesler D. Tackling COVID-19 with neutralizing monoclonal antibodies. Cell.

3. Yuan M, et al. Structural and functional ramifications of antigenic drift in recent SARS-CoV-2 variants. Science (New York, NY), (2021).

4. Volz E, et al. Evaluating the Effects of SARS-CoV-2 Spike Mutation D614G on Transmissibility and Pathogenicity. Cell 184, 64–75.e11 (2021).

5. Davies NG, et al. Estimated transmissibility and impact of SARS-CoV-2 lineage B.1.1.7 in England. Science (New York, NY) 372, (2021).

6. Tegally H, et al. Detection of a SARS-CoV-2 variant of concern in South Africa. Nature 592, 438–443 (2021).

7. Faria NR, et al. Genomics and epidemiology of the P.1 SARS-CoV-2 lineage in Manaus, Brazil. Science (New York, NY) 372, 815–821 (2021).

8. Cherian S, et al. Convergent evolution of SARS-CoV-2 spike mutations, L452R, E484Q and P681R, in the second wave of COVID-19 in Maharashtra, India. bioRxiv : the preprint server for biology, (2021).

9. Chen RE, et al. Resistance of SARS-CoV-2 variants to neutralization by monoclonal and serum-derived polyclonal antibodies. Nature medicine, (2021).

10. Hoffmann M, et al. SARS-CoV-2 variants B.1.351 and P.1 escape from neutralizing antibodies. Cell, (2021).

11. Hoffmann M, et al. SARS-CoV-2 variant B.1.617 is resistant to Bamlanivimab and evades antibodies induced by infection and vaccination. bioRxiv : the preprint server for biology, (2021).

12. Garcia-Beltran WF, et al. Multiple SARS-CoV-2 variants escape neutralization by vaccine-induced humoral immunity. Cell 184, 2372–2383.e2379 (2021).

13. Edara VV, et al. Infection- and vaccine-induced antibody binding and neutralization of the B.1.351 SARS-CoV-2 variant. Cell host & microbe 29, 516–521.e513 (2021).

14. Wu K, et al. Serum Neutralizing Activity Elicited by mRNA-1273 Vaccine. The New England journal of medicine 384, 1468–1470 (2021).

15. Wang Z, et al. mRNA vaccine-elicited antibodies to SARS-CoV-2 and circulating variants. Nature, (2021).

16. Chen X, et al. Disease severity dictates SARS-CoV-2-specific neutralizing antibody responses in COVID-19. Signal transduction and targeted therapy 5, 180 (2020).

17. Sokal A, et al. Maturation and persistence of the anti-SARS-CoV-2 memory B cell response. Cell, (2021).

18. Bao L, et al. The pathogenicity of SARS-CoV-2 in hACE2 transgenic mice. Nature 583, 830–833 (2020).

19. Sun SH, et al. A Mouse Model of SARS-CoV-2 Infection and Pathogenesis. Cell host & microbe 28, 124–133.e124 (2020).

20. Wrapp D, et al. Cryo-EM structure of the 2019-nCoV spike in the prefusion conformation. Science (New York, NY) 367, 1260–1263 (2020).

21. Hsieh CL, et al. Structure-based design of prefusion-stabilized SARS-CoV-2 spikes. Science (New York, NY) 369, 1501–1505 (2020).

22. Du S, et al. Structurally Resolved SARS-CoV-2 Antibody Shows High Efficacy in Severely Infected Hamsters and Provides a Potent Cocktail Pairing Strategy. Cell 183, 1013–1023 e1013 (2020).

23. Barnes CO, et al. SARS-CoV-2 neutralizing antibody structures inform therapeutic strategies. Nature 588, 682–687 (2020).

24. Benton DJ, et al. Receptor binding and priming of the spike protein of SARS-CoV-2 for membrane fusion. Nature 588, 327–330 (2020).

25. Lan J, et al. Structure of the SARS-CoV-2 spike receptor-binding domain bound to the ACE2 receptor. Nature 581, 215–220 (2020).

26. Pinto D, et al. Cross-neutralization of SARS-CoV-2 by a human monoclonal SARS-CoV antibody. Nature 583, 290–295 (2020).

27. Romero PE, et al. The Emergence of SARS-CoV-2 Variant Lambda (C.37) in South America. medRxiv : the preprint server for health sciences, (2021).

28. Toelzer C, et al. Free fatty acid binding pocket in the locked structure of SARS-CoV-2 spike protein. Science (New York, NY) 370, 725–730 (2020).

29. Zhang J, et al. Structural impact on SARS-CoV-2 spike protein by D614G substitution. Science (New York, NY) 372, 525–530 (2021).

30. Dejnirattisai W, et al. Antibody evasion by the P.1 strain of SARS-CoV-2. Cell 184, 2939–2954.e2939 (2021).

31. Wang P, et al. Increased resistance of SARS-CoV-2 variant P.1 to antibody neutralization. Cell host & microbe 29, 747–751.e744 (2021).

32. Lv Z, et al. Structural basis for neutralization of SARS-CoV-2 and SARS-CoV by a potent therapeutic antibody. Science (New York, NY) 369, 1505–1509 (2020).

33. Tortorici MA, et al. Ultrapotent human antibodies protect against SARS-CoV-2 challenge via multiple mechanisms. Science (New York, NY) 370, 950–957 (2020).

34. Huo J, et al. Neutralization of SARS-CoV-2 by Destruction of the Prefusion Spike. Cell host & microbe 28, 497 (2020).

35. Asarnow D, et al. Structural insight into SARS-CoV-2 neutralizing antibodies and modulation of syncytia. Cell 184, 3192–3204 e3116 (2021).

36. Yuan M, et al. A highly conserved cryptic epitope in the receptor binding domains of SARS-CoV-2 and SARS-CoV. Science (New York, NY) 368, 630–633 (2020).

37. Cohen J. South Africa Suspends Use of AstraZeneca’s COVID-19 Vaccine after It Fails to Clearly Stop Virus Variant. Science Magazine, February 7, 2021. https://www.sciencemag.org/news/2021/02/south-africa-suspendsuse-astrazenecas-covid-19-vaccine-after-it-fails-clearly-stop., (2021).

38. Wang GL, et al. Susceptibility of Circulating SARS-CoV-2 Variants to Neutralization. The New England journal of medicine, (2021).

39. Herper M, Branswell, H. New Data Shed Light on Efficacy of J&J’s Single-Dose Covid Vaccine. Stat, February 24, 2021. https://www.statnews/.com/2021/02/24/new-data-shed-light-on-efficacy-of-jjs-single-dose-vaccineagainst-covid-19/. (2021).

40. Xu K, et al. Epitope-based vaccine design yields fusion peptide-directed antibodies that neutralize diverse strains of HIV-1. Nature medicine 24, 857–867 (2018).

41. Kong R, et al. Fusion peptide of HIV-1 as a site of vulnerability to neutralizing antibody. Science (New York, NY) 352, 828–833 (2016).

42. Rappazzo CG, et al. Broad and potent activity against SARS-like viruses by an engineered human monoclonal antibody. Science (New York, NY) 371, 823–829 (2021).

43. Chen X, et al. Human monoclonal antibodies block the binding of SARS-CoV-2 spike protein to angiotensin converting enzyme 2 receptor. Cellular & molecular immunology 17, 647–649 (2020).

44. Smith K, et al. Rapid generation of fully human monoclonal antibodies specific to a vaccinating antigen. Nature protocols 4, 372–384 (2009).

45. Real-Fernández F, et al. Surface plasmon resonance-based methodology for anti-adalimumab antibody identification and kinetic characterization. Analytical and bioanalytical chemistry 407, 7477–7485 (2015).

46. Ju B, et al. Human neutralizing antibodies elicited by SARS-CoV-2 infection. Nature 584, 115–119 (2020).

47. Sun J, et al. Generation of a Broadly Useful Model for COVID-19 Pathogenesis, Vaccination, and Treatment. Cell 182, 734–743.e735 (2020).

48. Lei J, Frank J. Automated acquisition of cryo-electron micrographs for single particle reconstruction on an FEI Tecnai electron microscope. J Struct Biol 150, 69–80 (2005).

49. Zheng SQ, Palovcak E, Armache JP, Verba KA, Cheng Y, Agard DA. MotionCor2: anisotropic correction of beam-induced motion for improved cryo-electron microscopy. Nat Methods 14, 331–332 (2017).

50. Punjani A, Rubinstein JL, Fleet DJ, Brubaker MA. cryoSPARC: algorithms for rapid unsupervised cryo-EM structure determination. Nat Methods 14, 290–296 (2017).

51. Henderson R, et al. Outcome of the first electron microscopy validation task force meeting. Structure 20, 205–214 (2012).

52. Scheres SH, Chen S. Prevention of overfitting in cryo-EM structure determination. Nat Methods 9, 853–854 (2012).

53. Emsley P, Lohkamp B, Scott WG, Cowtan K. Features and development of Coot. Acta Crystallogr D Biol Crystallogr 66, 486–501 (2010).

54. Liebschner D, et al. Macromolecular structure determination using X-rays, neutrons and electrons: recent developments in Phenix. Acta Crystallogr D Struct Biol 75, 861–877 (2019).

55. Pettersen EF, et al. UCSF ChimeraX: Structure visualization for researchers, educators, and developers. Protein science : a publication of the Protein Society 30, 70–82 (2021).

